# Age-related differences in functional asymmetry during memory retrieval revisited: no evidence for contralateral over-activation or compensation

**DOI:** 10.1101/419739

**Authors:** James M. Roe, Didac Vidal-Piñeiro, Markus H. Sneve, Kristiina Kompus, Douglas N. Greve, Kristine B. Walhovd, Anders M. Fjell, René Westerhausen

## Abstract

Brain asymmetry is inherent to cognitive processing and seems to reflect processing efficiency. Lower frontal asymmetry is often observed in older adults during memory retrieval, yet it is unclear whether lower asymmetry implies an age-related increase in contralateral recruitment, whether less asymmetry reflects compensation, is limited to frontal regions, or predicts neurocognitive stability or decline. We assessed age-differences in asymmetry across the entire cerebral cortex, using fMRI data from 89 young and 76 older adults during successful retrieval, and surface-based methods that allowed direct homotopic comparison of activity between hemispheres. An extensive left-asymmetric network facilitated retrieval in both young and older adults, whereas diverse frontal and parietal regions exhibited lower asymmetry in older adults. However, lower asymmetry was not associated with age-related increases in contralateral recruitment, but primarily reflected either less deactivation in contralateral regions reliably signalling retrieval failure in the young, or lower recruitment of the dominant hemisphere—suggesting that functional deficits may drive lower asymmetry in older brains, not compensatory activity. Lower asymmetry neither predicted current memory performance, nor the extent of memory change across the preceding ∼8 years in older adults. Together, these findings are inconsistent with a compensation account for lower asymmetry during retrieval and aging.

## 1. Introduction

Episodic memory decline is a hallmark of neurocognitive aging (Park and Reuter-Lorenz 2009; Josefsson et al. 2012). Hence, understanding how alterations in brain functioning relate to aging and memory performance is a major research goal (Grady et al. 2006; Reuter-Lorenz and Park 2010a; Spreng et al. 2010; Grady 2012; Nyberg et al. 2012; Fjell et al. 2014). Lower functional asymmetry has been consistently observed in specific cortical regions in older relative to younger adults during memory tasks (Cabeza, 2002), and it has been proposed this may support optimal memory performance in older adults (Cabeza et al. 2002). However, whether older age is associated with anatomically widespread differences in asymmetry during memory processing, and the impact of age-related asymmetry differences upon memory performance, remain open questions. The purpose of the present study was to identify age-related differences in asymmetry across the entire cerebral cortex during memory retrieval, assess asymmetry relationships with current memory performance, and assess whether asymmetry may be a marker for memory preservation or decline over time in neurocognitive aging.

Functional asymmetry is thought to support processing efficiency (Hellige 1993; Ringo et al. 1994; Corballis 2009; Ocklenburg and Güntürkün 2012; Gotts et al. 2013). Previous evidence indicates that prefrontal asymmetry is lower in older adults during episodic memory retrieval tasks (Bäckman et al. 1997; Cabeza et al. 1997, 2002, 2004; Madden et al. 1999; Rossi et al. 2004), and such findings have been generalised within the Hemispheric Asymmetry Reduction in OLDer adults model (HAROLD). Although various theories have been proposed to account for this (Li et al. 2001; Cabeza 2002; Reuter-Lorenz 2002; Reuter-Lorenz and Park 2010a; Cabeza and Dennis 2012), most agree that the phenomenon reflects diminished processing efficiency at higher age, such that older brains are less able to rely on asymmetric processing strategies. Arguably the most influential (Cabeza et al. 2005; Reuter-Lorenz and Lustig 2005; Bishop et al. 2010; Grady 2012), the compensation view proposes that older adults recruit additional resources from the contralateral/subdominant hemisphere to cope with a declining ability of the dominant hemisphere to meet memory demands (Cabeza 2002; Cabeza et al. 2004; Cabeza and Dennis 2012). In line with this, most of the reported age-related differences in asymmetry during retrieval-related processing (Bäckman et al. 1997; Cabeza et al. 1997, 2002; Madden et al. 1999; Grady et al. 2002) and successful retrieval (Cabeza et al. 2004; Rossi et al. 2004) appear to show an apparent age-related increase in contralateral prefrontal recruitment, and have consequently been predominantly interpreted as compensatory (Cabeza 2002; Cabeza and Dennis 2012; Grady 2012; Wang and Cabeza 2017).

This interpretation is often reinforced when contralateral recruitment has a beneficial effect on memory performance, either across the group of older adults as a whole, or within low-performing older adults thought to rely more on compensation to maintain performance (Morcom and Johnson 2015; Cabeza et al. 2018). However, reported relationships between contralateral recruitment and cognitive function—including memory retrieval— remain highly inconsistent (Eyler et al. 2011; Grady 2012). Furthermore, high stability of individual differences in cognitive function across the lifespan (Deary et al. 2004; Hansen et al. 2014) challenges the assumption that low-performing older adults have undergone the most age-related decline (Rugg 2017). As a result, it is unclear whether lower asymmetry may be a marker for memory preservation or decline with advanced age.

Because the term compensation has typically been invoked to describe instances of enhanced brain activity in older adults (Cabeza et al. 2018), the compensation view of HAROLD arguably places greater emphasis (implicitly or otherwise) on contralateral over-recruitment in driving lower asymmetry effects in older brains. Yet, as noted by others (Cabeza 2002), lower asymmetry does not necessarily imply greater contralateral recruitment, because insufficient engagement of the ipsilateral/dominant hemisphere could also amount to less asymmetry. Thus, to better understand the role of asymmetry change in aging, one should evaluate asymmetry across large samples in a manner that comprehensively assesses each hemisphere’s contribution to the observed age-related differences in asymmetry.

In part due to methodological challenges in comparing activity between cerebral hemispheres that are anatomically asymmetric, most of the memory literature has focused on age-related differences in prefrontal asymmetry, and delineated target areas using a region-of-interest approach (Wilke and Lidzba 2007). Although often useful, such an approach limits the study of age-related differences in memory-asymmetries to specific brain regions (Berlingeri et al. 2013). Further, because hemispheric comparisons are usually informed by qualitative differences in the brain activation patterns of young and older adults, important questions have been overlooked, such as to what extent age-related differences in asymmetry are only partial (i.e. reflect a relative difference in asymmetry in regions that are nevertheless recruited asymmetrically in both age-groups; Westerhausen et al. 2014), or whether less asymmetry is also evident in regions that are recruited bilaterally (Cabeza and Dennis 2012). Additionally, because contrasts used to delineate retrieval-related activity have not always isolated activity associated with successful retrieval (Bäckman et al. 1997; Cabeza et al. 1997, 2002; Madden et al. 1999), it is unclear whether putative age-related over-recruitment of the contralateral hemisphere is indicative of higher memory success effects, or reduced negative success effects (i.e. lower retrieval failure effects). To address these questions, it is necessary to rely on an unbiased procedure that allows direct hemispheric comparisons on a more global scale. To our knowledge, the only existing study comparing whole-brain retrieval-asymmetry between young and older adults (Berlingeri et al. 2013) indicated that age-related asymmetry differences may be anatomically widespread, and that the underlying patterns are likely both complex and regionally dependent—thus underscoring the importance of conducting global asymmetry assessments.

Finally, although earlier research most often linked retrieval to a right-hemispheric prefrontal specialization (Tulving et al. 1994; Cabeza et al. 1997; Habib et al. 2003), an abundance of more recent evidence suggests successful retrieval is associated with predominantly left-lateralized cortical activation, also in prefrontal cortex (Morcom et al. 2007; McDermott et al. 2009; Spaniol et al. 2009; Kim 2010, 2016; de Chastelaine et al. 2016; Vidal-Piñeiro et al. 2017). Meta-analytic evidence suggests that left hemisphere effects may be particularly prominent in lateral frontal and parietal regions during the retrieval of events in laboratory-based tasks (McDermott et al. 2009; Spaniol et al. 2009; Spreng et al. 2010; Kim 2016), possibly reflecting the relevance of verbal or verbalizable components (Kim 2016). Hence, left-lateralization may represent a prominent feature of retrieval networks likely to be engaged during the type of memory task employed here (Vidal-Piñeiro et al. 2017). Given the decline in source memory as we age (Anderson et al. 2008; Old and Naveh-Benjamin 2008; Cansino et al. 2013, 2019; Gorbach et al. 2017; Langnes et al. 2018), the expected relationship between memory ability and isolated memory success activity (Rugg 2017), and putative reductions in retrieval-asymmetry in older adults, source memory retrieval represents an ideal task to assess age-related differences in asymmetry and their relation to neurocognitive aging.

Here, we directly investigated age-related differences in functional asymmetry across the entire cerebral cortex during an item-source associative memory retrieval task, using fMRI data from a large sample of participants. We delineated asymmetry via surface-based analyses that directly contrast homotopic activity between the cortical hemispheres, applying an unbiased symmetrical template (Greve et al. 2013; Maingault et al. 2015; Tobyne et al. 2016; Greve and Fischl 2017). Since we were interested in age-related differences in retrieval-asymmetry, we contrasted activity elicited during the presentation of items recalled with an accurate source memory judgement with activity elicited during the presentation of items not remembered (i.e. source versus miss). Although contrast choice is a matter of debate (see discussion), we chose this over a source versus item-memory contrast to avoid confounding age-related differences in retrieval-related activity with age-related differences in activity associated with other cognitive processes known to be affected by age—such as post-retrieval monitoring (McDonough and Gallo 2013; McDonough et al. 2013; Mitchell et al. 2013; de Chastelaine et al. 2016), the demands for which are thought to be greater during the retrieval of weaker (i.e. item) versus stronger (i.e. source) memory signals (Henson et al. 1999; Achim and Lepage 2005; Kim 2010; de Chastelaine et al. 2016; Wang et al. 2016).

The main objectives of the study were two-fold. First, to investigate support for the compensation view of HAROLD during memory retrieval, we aimed to 1) examine whether global differences in asymmetry are apparent in older relative to younger adults during retrieval, 2) assess whether the pattern of lower asymmetry is indicative of greater contralateral recruitment in older adults relative to the young, and 3) test whether lower asymmetry predicts current memory performance. The conditions to interpret the data within a neural compensation framework were pre-established; in line with recently published guidelines (see Cabeza et al. 2018), we resolved that support for neural compensation would require 1) evidence of enhanced brain activity in older adults in 2) the form of greater contralateral recruitment relative to the young, and 3) evidence that lower asymmetry has a beneficial effect on current memory performance.

Second, we aimed to test whether the extent of asymmetry—as a putative proxy for processing efficiency—may be a marker for memory preservation or decline in neurocognitive aging, using multi-timepoint longitudinal data obtained over the ∼8-year period prior to scanning in older participants.

## 2. Materials & methods

### 2.1 Participants

Participants were drawn from the third waves of ongoing longitudinal projects coordinated by the Center for Lifespan Changes in Brain and Cognition (Department of Psychology, University of Oslo) for which task fMRI data were available: *Cognition and Plasticity through the Lifespan* (Sneve et al. 2015; Vidal-Piñeiro et al. 2017) and *Neurocognitive Development* (Tamnes et al. 2018). The sample consisted of 165 participants, of which 89 were young aged 18-39 (56 female, mean age = 26.8 [standard deviation (SD) = 5.0], age range = 18.5 - 38.9 years) and 76 were older aged 60 and above (40 female, mean age = 67.9 [SD = 5.2], age range = 60.2 – 80.8 years). Additionally, for 54 of the older participants, longitudinal cognitive data spanning back ∼8 years were available. 51 of these had data from all three waves, whereas 3 participants were recruited at wave 1 and again at wave 3 (mean years between wave 1 and 3 = 7.7 [SD = 0.9]).

All participants were fluent in Norwegian, reported normal or corrected-to-normal hearing and vision, and no motor difficulties. All reported no history of neurological or psychiatric disorders, and none were taking any medication known to affect CNS functioning. Participants were required to be right-handed, have a score of <16 on the Beck Depression Inventory (Beck et al. 1961), and score a normal IQ (≥ 85) on the Wechsler Abbreviated Scale of Intelligence (WASI; Wechsler 1999), although fMRI data for eight young participants with missing neuropsychological data was also included. Additionally, older participants were required to score ≥ 25 on the Mini Mental Status Exam (MMSE; Folstein et al. 1975), a normative score of t ≥ 30 on the California Verbal Learning Test (CVLT; Delis et al. 1987) encoding and 30 minute delayed recall, and to have recalled ≥ 10% of trials with source memory on the fMRI task (see Supplementary Table 1 for sample descriptives and neuropsychological data). For inclusion in longitudinal analyses, CVLT data from the first wave of data collection were required to meet the same criteria. All provided written informed consent, and the study received approval from the Regional Ethical Committee of South Norway. Participants were paid for their participation.

### 2.2 Experimental Design

The experiment consisted of an incidental source memory encoding task, and a surprise memory test after ∼90 minutes. Both encoding and retrieval phases were performed in the MRI scanner. At encoding, participants were asked to decide whether a particular action (either “can you lift it?” or “can you eat it?”) could be performed on an item, and were later tested on their memory for these item-action associations under a test phase occurring ∼1.5 hours later. The entire experiment consisted of 2 incidental encoding runs and 4 test runs, and has been thoroughly described elsewhere (Sneve et al. 2015; Vidal-Piñeiro et al. 2017, 2018). Only fMRI data from the test phase is used here. Briefly, encoding stimuli consisted of 100 monochromatic line drawings (50 per run) depicting everyday objects and items (Fig. 1; left). At encoding, each question (lift/eat; aurally presented by a female voice in the Norwegian language) was presented 25 times per run in a pseudorandom order, and one second after question onset, an image of an item was presented on screen together with a response indicator that instructed participants which button to press to respond with a ‘yes’ [the item can be lifted/eaten] or ‘no’ [the item cannot be lifted/eaten]. Button-response mapping was counterbalanced and participants had the 2 seconds (s) stimulus duration to respond.

**Figure 1.**
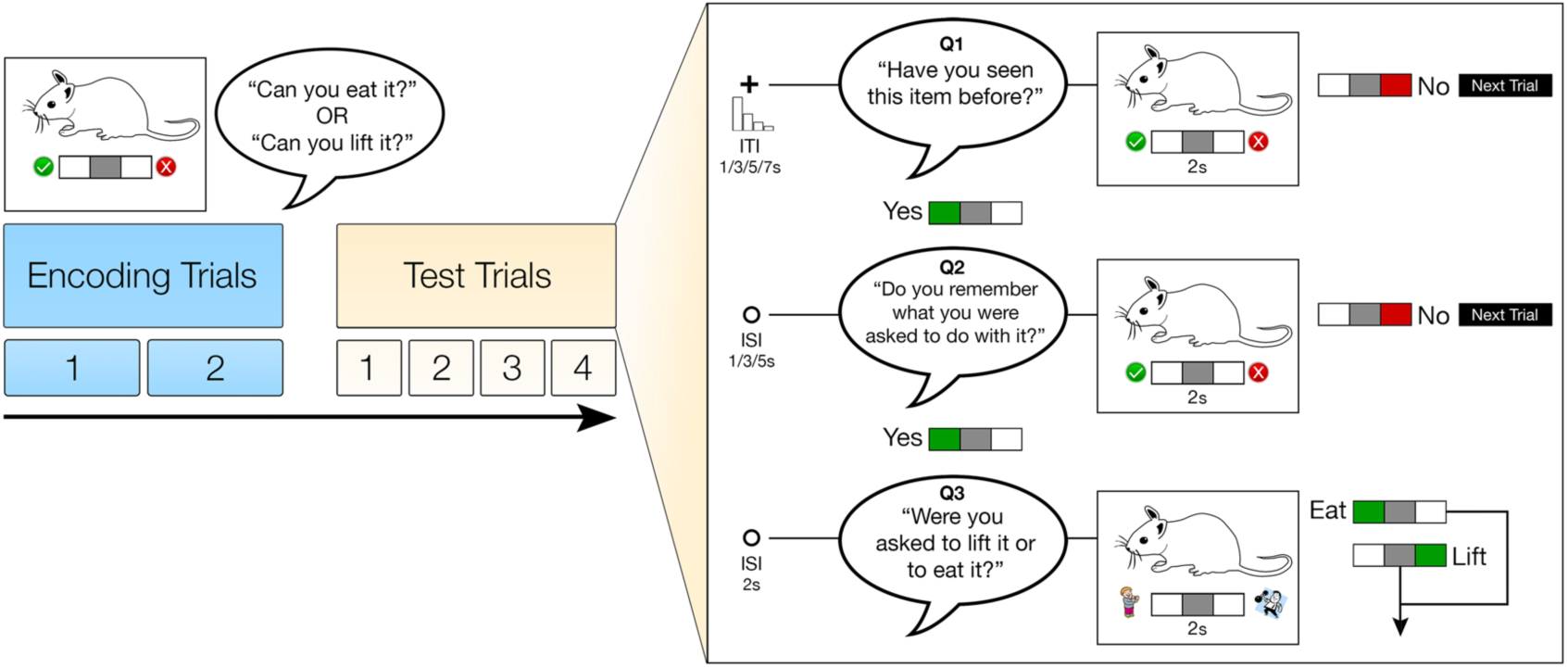
Experimental paradigm. Participants underwent 100 incidental encoding trials (2 fMRI runs) and were tested for memory for item + action (lift/eat) associations (4 fMRI runs) ∼1.5 hours later. At test, old items were randomly mixed with 100 new items. The fMRI retrieval paradigm consisted of a three-question (Q1-Q3) procedure (2s duration per Q) where progression to the next was contingent upon a ‘Yes’ response to the previous. Q3 consisted of a two-alternative forced choice between actions (lift/eat). The trial ended on ‘No’ responses to Q1 or Q2. Response cues were present on screen, and button response mapping was counterbalanced across participants. Crucially, only activity corresponding to initial stimulus presentation (Q1) was used in all analyses. ITI/ISI = Inter-trial/-stimulus interval, respectively.

Approximately 90 minutes later participants received a surprise memory test, where each of the 100 ‘old’ items were tested intermixed with 100 new ‘foils’ in an event-related fMRI paradigm. Each of the 4 test runs consisted of 25 ‘old’ mixed with 25 ‘new’ items, presented in a pseudorandom order. All runs started and ended with a 11s baseline recording period in which a black central fixation-cross was presented against a white background. An additional baseline period was presented once in the middle of each run. A test trial began with the same female voice asking the Norwegian equivalent of: “Have you seen this item before?” This constituted the first question (Q1) in a three-question procedure (see Fig. 1), where progression to the next question was contingent upon a ‘yes’ response to the previous. 1s after question onset, an item appeared on screen together with the response indicator. For Q1, participants were to respond either ‘yes’ [I saw the item during the encoding phase] or ‘no’ [I did not see the item during the encoding phase] within the 2s duration the stimulus remained on screen. A ‘no’ response (or a missed response) to Q1 ended the trial. A fixation-cross remained on screen throughout the following inter-trial interval (ITI), which lasted for 1-7s (exponential distribution over four discrete intervals; ITI order tentatively optimised using optseq2 (http://surfer.nmr.mgh.harvard.edu/optseq/)). If the participant believed they remembered seeing the item (responded ‘yes’), the trial proceeded to the second question (Q2): “Do you remember what you were asked to do with it?”. A ‘no’ response to Q2 ended the trial. A ‘yes’ response indicated that the participant remembered the associated action, and prompted the final two-alternative forced-choice control question (Q3): “Were you asked to lift it or eat it?” [I was asked whether the item could be lifted/eaten during the encoding phase]. Q2 was thus included to dissuade per-item guessing behaviour prior to the control question. Memory was then determined for each old item according to each participant’s response at test.

### 2.3 Behavioural analysis

Test trial responses to old items were behaviourally classified as follows: (1) *source memory* (‘yes’ response to Q1 and Q2, and correct response to Q3); (2) *item-only memory* (correct ‘yes’ response to Q1 and either a ‘no’ response to Q2 or incorrect response to Q3); or (3) *miss* (incorrect ‘no’ response to Question 1). New items were classified either as correct rejections or false alarms based on the response at Q1. Although the three-question procedure was optimised to disentangle true source memories from item-only and missed memories, it nevertheless remained possible to achieve 100% source memory given straight “yes” responses to all three questions on all trials. Thus, the raw percentage of source memory hits was corrected for by subtracting the number of incorrect responses at Q3 (source memory – incorrect source memory). This correction tentatively controls for individual differences in false source memories, threshold criteria at Q2, or guessing behaviour that affects raw source memory estimates (Vidal-Piñeiro et al. 2017). All behavioural analyses were performed in R (https://www.r-project.org; v. 3.3.2), and Bonferroni correction was applied where appropriate, as indicated.

### 2.4 MRI acquisition and equipment

All functional and anatomical images were acquired on a Siemens Skyra 3T scanner using a Siemens 24-channel head coil (Siemens Medial Solutions, Germany) at Rikshospitalet, Oslo University Hospital. Functional data were acquired using a BOLD sensitive T2*-weighted EPI sequence, with equivalent acquisition parameters across all fMRI runs; each EPI volume consisted of 43 transversally-oriented slices taken using interleaved acquisition (no gap) acquired with the following parameters: TR = 2390ms; TE = 30ms; flip angle = 90°; voxel size = 3 × 3 × 3mm, FOV = 224 × 224mm; GRAPPA acceleration factor = 2). At the onset of each run, 3 dummy volumes were taken to account for a potentially imperfect flip angle due to T1 saturation effects, and were subsequently discarded. The mean number of volumes per retrieval run was 208.

Anatomical T1-weighted magnetization prepared gradient echo (MPRAGE) scans composed of 176 sagittally-oriented slices were acquired using a turbo-field echo pulse sequence with the following parameters: TR = 2300ms, TE = 2.98ms, flip angle = 8°, voxel size = 1 × 1 × 1mm, FOV = 256 × 256mm. Additionally, a standard double gradient-echo field map sequence was acquired for distortion of the echo planar images. Stimuli were presented on a NNL 32” LCD screen (resolution = 1920 × 1080px; NordicNeuroLab, Norway) viewed through a mirror mounted onto the scanner head-coil (176cm from mirror to screen). Auditory stimuli were presented via the scanner intercom to participant headphones, and responses were given using the ResponseGrip system (NordicNeuroLab). All stimuli were presented using E-Prime 2.0 stimulus presentation software (Psychology Software Tools, PA), and stimulus presentation was synchronised with MRI image acquisition via a NNL SyncBox (NordicNeuroLab).

### 2.5 MRI preprocessing

Cortical reconstruction and volumetric segmentation of the T1-weighted anatomical images was performed using FreeSurfer 5.3 (http://surfer.nmr.mgh.harvard.edu/; Dale et al. 1999; Fischl et al. 1999; Fischl and Dale 2000). Briefly, this automated pipeline included motion correction, removal of non-brain tissue, Talairach transformation, intensity normalisation, demarcation of grey/white and grey/CSF boundaries at the locations where the greatest intensity shifts reflect the transition between tissue classes, and inflation of the cortical surface (Fischl, Sereno, and Dale 1999b; Fischl, Sereno, Tootell, et al. 1999).

fMRI data was preprocessed using a combination of FSL (http://fsl.fmrib.ox.ac.uk/fsl/fslwiki) and FreeSurfer Functional Analysis Stream (FSFAST; https://surfer.nmr.mgh.harvard.edu/fswiki/FsFast). First, using FSL’s fMRI Expert Analysis Tool functional images were corrected for distortions caused by B0 inhomogeneities in EPI scans, slice-timing corrected to the middle TR in the volume, high-pass filtered with a cutoff of 0.01Hz, and slightly smoothed in volume space (5mm FWHM) as is recommended prior to running ICA-based denoising methods (Griffanti et al. 2014; Salimi-Khorshidi et al. 2014). Next, FSĹs Multivariate Exploratory Linear Optimized Decomposition into Independent Components (MELODIC) was used together with FMRIB’s ICA-based Xnoiseifier (FIX) tool, applying a custom training file (28 participants aged 18-80) to auto-classify components into “good” and “bad” components, and remove bad components from the 4D data (Salimi-Khorshidi et al. 2014). During FIX, the unique variance associated with 6 motion parameters (and the derivative up to the second-polynomial; i.e. 18 parameters) was also non-aggressively regressed out of the 4D data. Such ICA-based procedures for denoising fMRI data have been shown to effectively reduce motion-induced variability, outperforming methods based on removing motion spikes (Pruim et al. 2015). Functional datasets were then intensity normalised and projected onto the cortical surface (i.e. vertex space) using FreeSurfer routines. This involved resampling of the preprocessed functional data to the reconstructed cortical hemispheres of each individual subject, using robust boundary-based registration algorithms that bring functional images into alignment with the reference anatomical image by maximizing the intensity gradient across tissue boundaries (Greve and Fischl 2009). Functional data was then smoothed in 2D surface space (8mm FWHM) prior to first-level statistical analyses. Such surface-based approaches to analyzing fMRI data are considered beneficial compared to volume-based approaches because they adhere to the inherent geometry of the cortical sheet (Fischl 2012; Glasser et al. 2016; Coalson et al. 2018).

### 2.6 fMRI analysis

#### 2.6.1 First-level fMRI analysis

Each of the 100 ‘old’ items was assigned to a condition/regressor based on the memory condition classified at test. A first-level GLM was set up modeling the onset times and stimulus durations (2s) for each retrieval regressor through convolution with a double-gamma canonical hemodynamic response function. The task regressors-of-interest were: *source memory* (item correctly remembered with associated action) and *miss* trials (item not remembered). Each regressor-of-interest modelled the onsets and durations corresponding to the initial stimulus presentation within the three-question procedure (i.e. Q1). Crucially, then, only activity elicited at Q1 was used in all analyses. A set of nuisance regressors were included to account for BOLD variance of non-interest, and included regressors for trials classified as item-only memory (item correctly remembered/recognised without associated action) and trials where stimuli could not be classified into a memory condition (due to failure to respond at any question). Additional regressors were included to model the BOLD response at Q2 and Q3, false alarms and correctly rejected new items. The temporal derivatives of all regressors were included to improve model fit, as well as a set of polynomial nuisance regressors (up to the order of 2), and temporal autocorrelations in the model residuals were corrected using pre-whitening. For each individual, a contrast of parameter estimates was computed for further analysis, consisting of *source memory v. miss* conditions (*retrieval-success* contrast).

#### 2.6.2 Asymmetry approach

The first-level analyses were performed on the reconstructed cortical left and right hemispheres of each participant, and thus respected the individual cortical geometry of each subject. We then leveraged the symmetrical template that exists in FreeSurfer (fsaverage_sym) to apply reregistration between the left and right cortical surfaces of each individual, and the *left* hemisphere of the symmetrical template. This allowed mapping of the first-level data from both hemispheres into a common analysis space. Crucially, given the symmetry of the template, the mapping was not biased to the cortical geometry of either hemisphere (Greve et al. 2013). See Figure 2 for a schematic illustration of the symmetrical registration (note the ‘flipping’ of right hemisphere data to the left symmetrical surface; bottom right). Next, the contrast maps in symmetrical space were fed into second-level group analyses.

**Figure 2.**
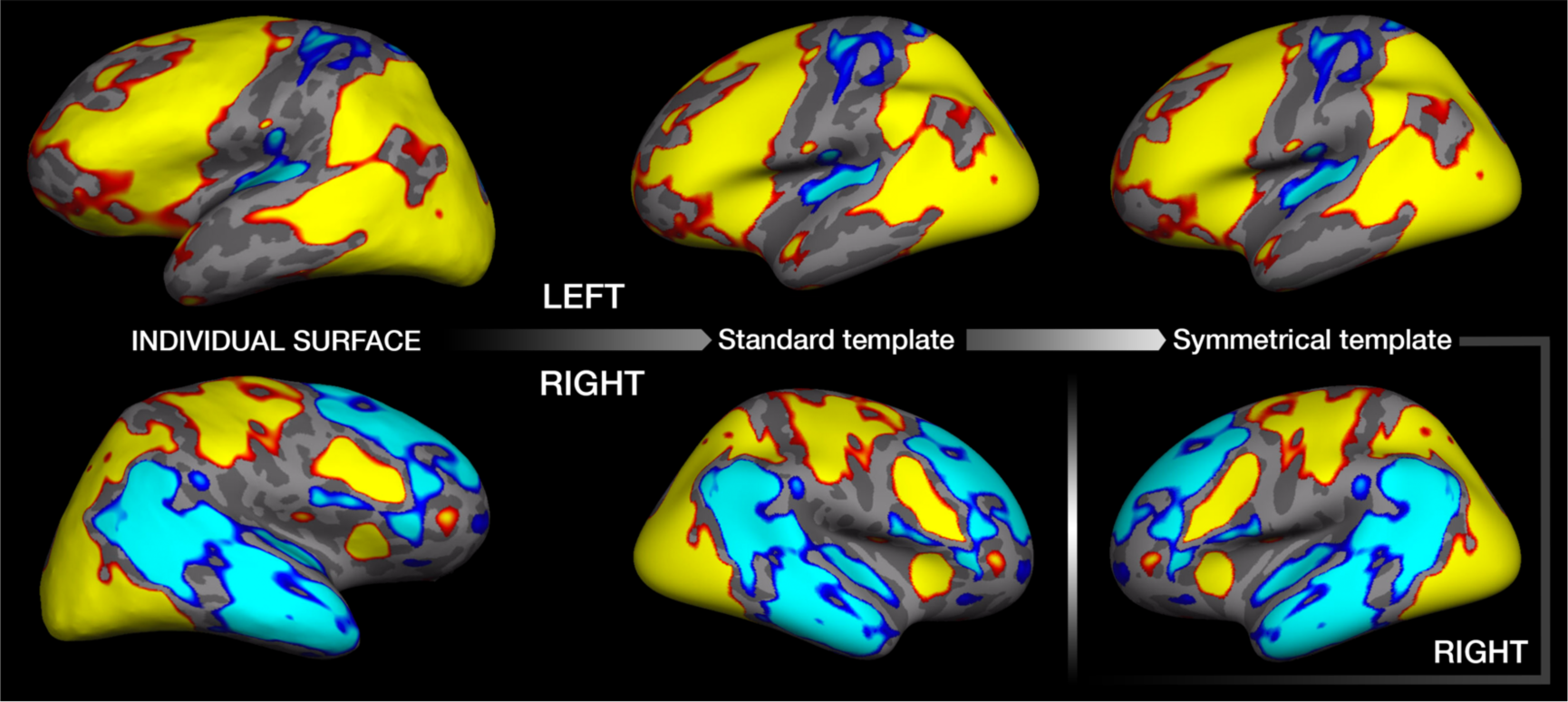
Single subject reregistration example. First level analyses were performed on the individual’s reconstructed cortical surface (left panel) and the results were subsequently reregistered to the standard symmetrical template provided by FreeSurfer (fsaverage_sym; right panel) prior to group-level analyses. Data corresponding to the left and right hemispheres are shown in the top and bottom panels, respectively. For visual comparison only, data is also shown reregistered to the non-symmetrical standard template (fsaverage; middle panel), although at no point was this used during processing. Note the ‘flipped’ right hemisphere data after reregistration to the left symmetrical template (bottom right panel). Data is shown for visualisation purposes only.

#### 2.6.3 Conventional fMRI analysis

Firstly, we performed vertex-wise group analyses following a conventional approach—that is, analyses were performed separately for each hemisphere—but within the symmetrical space. Group-level estimates for mean BOLD activity were computed on the cortical surface for 1) *Left* hemisphere (LH) retrieval-success activity in *Young*, 2) *Right* hemisphere (RH) retrieval-success activity in *Young*, 3) *LH* retrieval-success activity in *Old*, and 4) *RH* retrieval-success activity in *Old*. Ordinary least squares GLM analysis was performed for each, controlling for age and corrected source memory performance.

#### 2.6.4 fMRI asymmetry analysis

Having brought the data from each hemisphere into the symmetrical analysis space, age-related differences in asymmetry between young and older adults could be directly assessed via whole-cortical comparisons of homotopic activity on a vertex-wise basis. Here, individual cortical maps were fed into a mixed-effect 2 (Age-Group: young vs. old) × 2 (Hemisphere: left vs. right) ANOVA design, where group differences in brain activation asymmetry were tested with the Age-Group × Hemisphere interaction.

To better delineate the patterns of activity from the resulting interaction effects, the resulting clusters were split according to whether both hemispheres showed positive memory effects, negative memory effects, or whether the direction of the effect differed between the hemispheres. This approach allowed us to avoid averaging activity across functionally heterogeneous clusters. Here, we performed an interhemispheric conjunction analysis on the resulting clusters from the Age-Group × Hemisphere analysis (following correction and statistical thresholding; see below). For each hemisphere we computed the mean young and old *source memory v. miss* simple effect size map, binarized the maps on positive or negative values, and performed whole-cortical left × right multiplications of the binary maps to identify regions that on average exhibited positive effects in both hemispheres, negative effects in both hemispheres, or differential effects across hemispheres (i.e. positive in one and negative in the other). This approach yielded 9 functionally distinct ROI’s (>25 vertices). Importantly, as the ROI identification was based on the mean activation across age-groups, the signal change within ROI’s could be meaningfully compared between groups. Signal change was then extracted after averaging across all vertices in each ROI per participant, to describe the patterns underlying the Age-Group × Hemisphere interaction in post-hoc analyses.

Further analyses performed on the cortical surface investigated the separate main effects of Hemisphere and Age-Group in the mixed-effect design. The effect of hemisphere within both young and older adults was further explored via separate paired post-hoc *t*-tests (*left v. right*). This was performed to assess the degree of overlap in cortical activation patterns showing an asymmetry effect between young and old age-groups. Dice Similarity Coefficient was employed to compute the overlap (Crum et al. 2006).

Finally, since the main effect of Hemisphere does not differentiate between sites of absolute (i.e. significant effects only in one hemisphere) and relative asymmetry (i.e. significant effects in both hemispheres but significantly greater in one; Westerhausen et al. 2014), we performed post-hoc analyses to isolate absolute from relative asymmetry effects. Here, the mean RH activation map was calculated across age-groups, and a conjunction analysis was performed between the RH significance map and the significance map for the main effect of Hemisphere where the effects were positive (i.e. showed significantly greater effects in LH). The resulting map revealed cortical regions showing relative asymmetry, where both left and right homotopic cortex exhibited significant memory effects but where the effects in LH were still significantly greater.

For all cortical analyses, statistical significance was tested on a vertex-wise basis. The resulting maps were then corrected for multiple comparisons using a 2-step cluster-based approach, following recent recommendations (Eklund et al. 2016, 2018). First, a cluster-forming threshold was applied at p < 0.01, and clusters were tested through non-parametric permutation inference across 10,000 iterations (PALM; http://fsl.fmrib.ox.ac.uk/fsl/fslwiki/PALM; Winkler et al. 2014). Cluster significance was then considered at a family-wise error-corrected level of *p* < 0.05 (two-sided).

#### 2.6.5 Asymmetry relationships with current memory performance

Next, we tested whether the magnitude of asymmetry was associated with source memory performance on the fMRI task. For each participant and ROI, we computed an asymmetry metric by subtracting the right from the left signal change (LH - RH). Since ROI analyses revealed consistently higher activation in the left hemisphere relative to the right, higher values corresponded to a greater leftward asymmetry. In ROI’s that were positively activated in LH and deactivated in RH, this asymmetry index reflected the span between positive LH and negative RH activation. For each of the 9 ROI’s, we tested the relationship between asymmetry and memory via ANCOVA’s with Age-group, Source memory, and Age-group × Source memory as factors (age, sex controlled). In addition, in line with the current opinion that compensation is not necessarily expected to be prevalent in older high-performers, we post-hoc tested the view that compensatory brain responses may be most prominent in older participants with the greatest need (Cabeza and Dennis 2012; Morcom and Johnson 2015; Cabeza et al. 2018). Here, we median split older adults into low and high source memory performance-groups, and post-hoc tested the Group × Source memory interaction via ANCOVA’s (Group and Source memory as factors; age, sex controlled) to determine whether asymmetry-memory relationships differed by performance-group.

#### 2.6.6 Asymmetry relationships with longitudinal memory change

We next performed longitudinal analyses using a subset of older participants (N=54) to assess whether the extent of functional asymmetry in ROI’s showing age-related differences in asymmetry could be predicted by the degree of memory change exhibited across the preceding ∼8 years. Longitudinal memory change was estimated using data from multiple timepoints (Tp’s) in a memory task where performance was highly correlated with memory on our fMRI task (CVLT delayed recall). Specifically, 51 older adults had data from three-timepoints (corresponding to all three project waves), whereas 3 had two-timepoint data (recruited at wave 1 and again at wave 3). A three-step procedure was employed to estimate memory change: 1) we regressed each participants’ CVLT delayed recall scores against time to estimate an intercept and slope that represented baseline memory performance and memory change, respectively; 2) the slopes of all participants were modeled as a linear function of chronological age at the final timepoint (i.e. the third wave), partialling out the effects of sex (to account for differences in memory rates-of-change; Josefsson et al. 2012), and baseline performance at Tp1; 3) the residuals for the slope of each participant relative to this model were then calculated. This resulted in a continuous measure reflecting the deviation in predicted longitudinal memory change relative to each participants’ age at their final timepoint, corrected for baseline memory performance and sex. The residuals from the model were considered as a proxy for memory change. Linear regressions between asymmetry (see previous section) and memory change were then computed for each of the 9 ROI’s.

Finally, we computed the estimated effect size for memory change upon asymmetry that would be detectable by our current sample size via a posteriori sensitivity analysis using G*power 3.1 (α = 0.05, β = 0.80; Faul et al. 2007).

## 3. Results

### 3.1 Behavioural results

Following correction for the percentage of incorrect source memories (source memory hits minus incorrect source memories; see Table 1 to compare with raw source memory hits), young participants remembered on average 48.6% (± SD 13.5%) trials with source memory, whereas older participants remembered on average 28.4% (± 17.5%) trials with source. Source memory was significantly above chance level (0 post-correction) in both young (t(88)= 34.0, *p* < 10^-16^) and older age-groups (t(75) = 14.1, *p* < 10^-16^), and older participants remembered significantly fewer trials with source than the young (t(163) = −8.3, *p* = 10^-14^, Cohen’s d (*d*) *= -*1.3). For item-only memories (remembered without associated action), the difference between young (18.8% ± 6.6%) and older (21.2% ± 9.0%) participants was not significant (t(163) = 1.9, *p* = 0.06), and the number of forgotten items was also not significant between young (M = 22.3% ± 9.7%) and older adults (M = 25.1% ± 11.3%; t(163) = 1.7, *p* = 0.10). Additionally, the percentage of incorrect source memories (incorrect responses to Q3 for ‘old’ items) was significantly greater in older participants (M = 14.8% ± 7.7%) relative to young (M = 5.8% ± 5.2%; t(163) = 8.8, *p* = 10^-15^*, d =* 1.4). The number of trials per memory condition regressor was: 54.2 and 43.2 (source trials; young and old adults, respectively); 22.3 and 25.1 (miss trials; young and old adults, respectively). See Table 1 for full behavioural statistics.

**Table 1.**
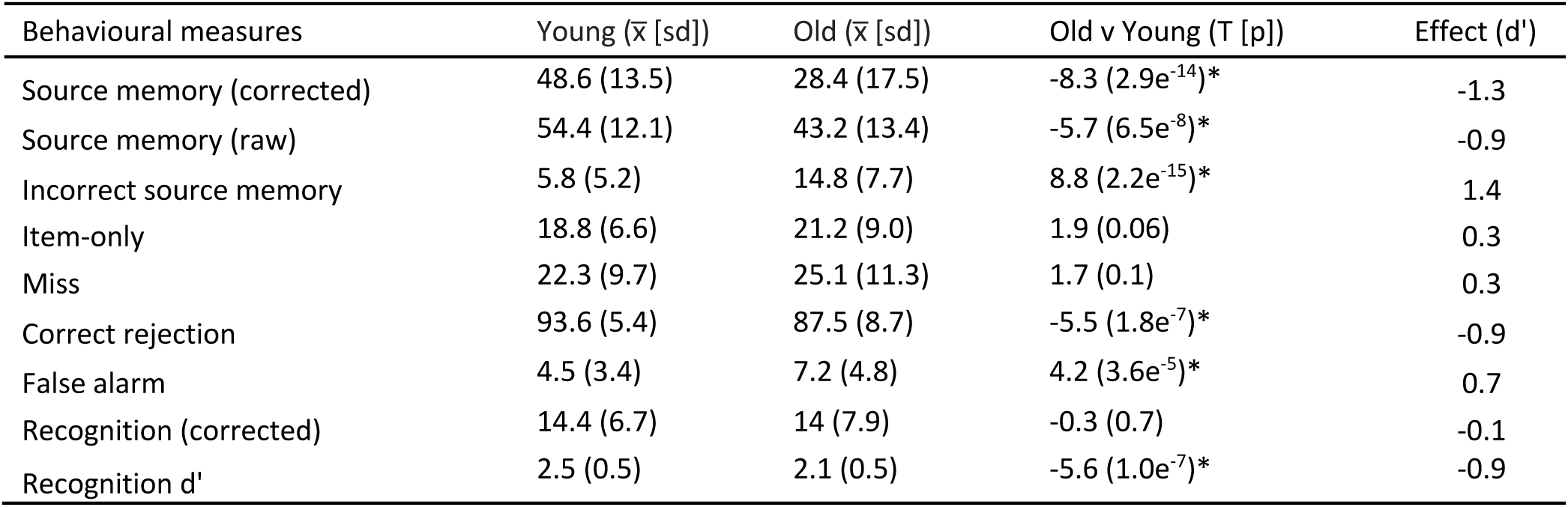
fMRI task behaviour.

| Behavioural measures | Young ($\bar{x}$ [sd]) | Old ($\bar{x}$ [sd]) | Old v Young ($T$ [$p$]) | Effect ($d'$) |
| --- | --- | --- | --- | --- |
| Source memory (corrected) | 48.6 (13.5) | 28.4 (17.5) | -8.3 ($2.9e^{-14}$)* | -1.3 |
| Source memory (raw) | 54.4 (12.1) | 43.2 (13.4) | -5.7 ($6.5e^{-8}$)* | -0.9 |
| Incorrect source memory | 5.8 (5.2) | 14.8 (7.7) | 8.8 ($2.2e^{-15}$)* | 1.4 |
| Item-only | 18.8 (6.6) | 21.2 (9.0) | 1.9 (0.06) | 0.3 |
| Miss | 22.3 (9.7) | 25.1 (11.3) | 1.7 (0.1) | 0.3 |
| Correct rejection | 93.6 (5.4) | 87.5 (8.7) | -5.5 ($1.8e^{-7}$)* | -0.9 |
| False alarm | 4.5 (3.4) | 7.2 (4.8) | 4.2 ($3.6e^{-5}$)* | 0.7 |
| Recognition (corrected) | 14.4 (6.7) | 14 (7.9) | -0.3 (0.7) | -0.1 |
| Recognition $d'$ | 2.5 (0.5) | 2.1 (0.5) | -5.6 ($1.0e^{-7}$)* | -0.9 |

Main behavioural measures from the fMRI memory task and results from *Old v. Young t*-tests for each measure (*Bonferroni corrected at *p* < [0.05/9] 0.006). Source memory (corrected) = Source memory (raw) – Incorrect source memory; Recognition (corrected) = Item-only – False alarms.

### 3.2 Conventional fMRI analysis: Exploring hemispheric effects irrespective of asymmetry

Vertex-wise analyses of the BOLD response associated with retrieval-success (*source v. miss*) were first conducted separately for each hemisphere and age-group, following a conventional approach. Significant activation was observed in widespread cortical regions of the LH in both young and older age-groups (Fig. 3; top row). Specifically, lateral regions of LH activation were observed encompassing large portions of frontal, middle temporal, and inferior and superior parietal cortex, whereas left medial sites included occipital, posteromedial parietal and superior frontal regions. Similar activation maps were obtained in the RH (Fig. 3; bottom row), albeit to a seemingly lesser degree, particularly in frontal and temporal regions. Dice similarity coefficient (DSC) revealed that cortical regions exhibiting significant positive retrieval-success effects were highly similar between young and older participants in both LH (DSC = 0.89) and RH (DSC = 0.79). In contrast, cortical regions exhibiting significant negative retrieval-success effects were only evident in young participants. These negative memory effects were encompassed within regions including bilateral supramarginal gyrus, superior temporal and insular cortices.

**Figure 3.**
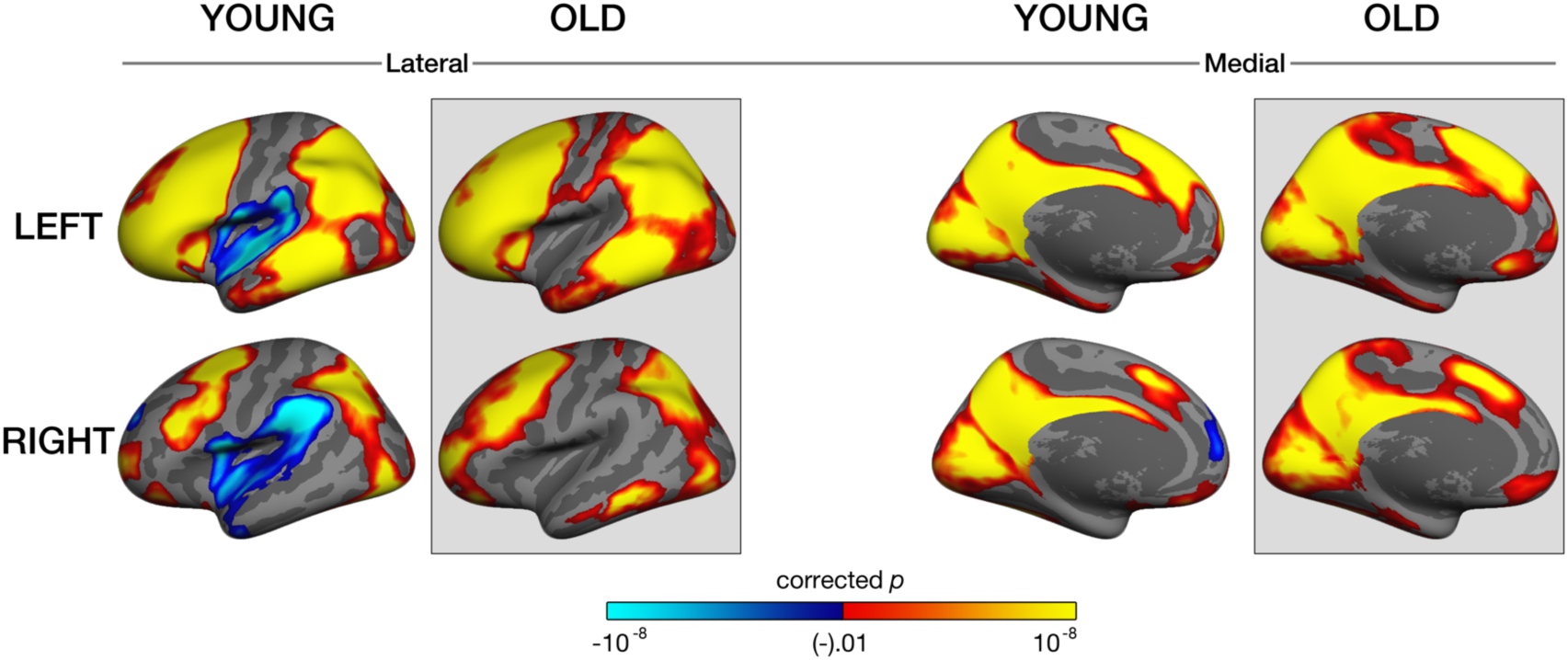
Results from conventional fMRI analyses performed separately by hemisphere for young and older adults (clusters surviving FWE multiple comparison correction; vertex-wise *p* < 0.01; cluster-based *p* < 0.05). All group analyses were performed within the symmetrical space (the left symmetrical surface). Note that the data from both left (top panel) and right (bottom panel) hemispheres is registered to the left symmetrical surface.

### 3.3 fMRI Asymmetry analysis

Next, we directly tested for differences in functional asymmetry across the whole cortex between young and older adults. A mixed-effect ANOVA revealed three clusters showing a significant Age-Group × Hemisphere interaction, indicating group differences in brain asymmetry (Fig. 4A). The significant clusters were localised in the lateral frontal, the supramarginal and the superior parietal cortices. An interhemispheric conjunction analysis revealed that these clusters consisted of 9 functionally distinct regions (Fig. 4B), which constituted the ROI’s for post-hoc analyses. Of these, 4 showed positive effects in both hemispheres (middle frontal, precentral, superior parietal and superior supramarginal cortex), 4 showed differential effects across hemispheres (i.e. positive in one and negative in the other; pars triangularis and opercularis of the inferior frontal gyrus, and two regions in supramarginal gyrus), and 1 showed negative effects in both hemispheres (anterior supramarginal gyrus). Significant age-related differences in asymmetry remained after partialling out between-subject differences in source memory performance and sex in all except one ROI (precentral gyrus).

**Figure 4.**
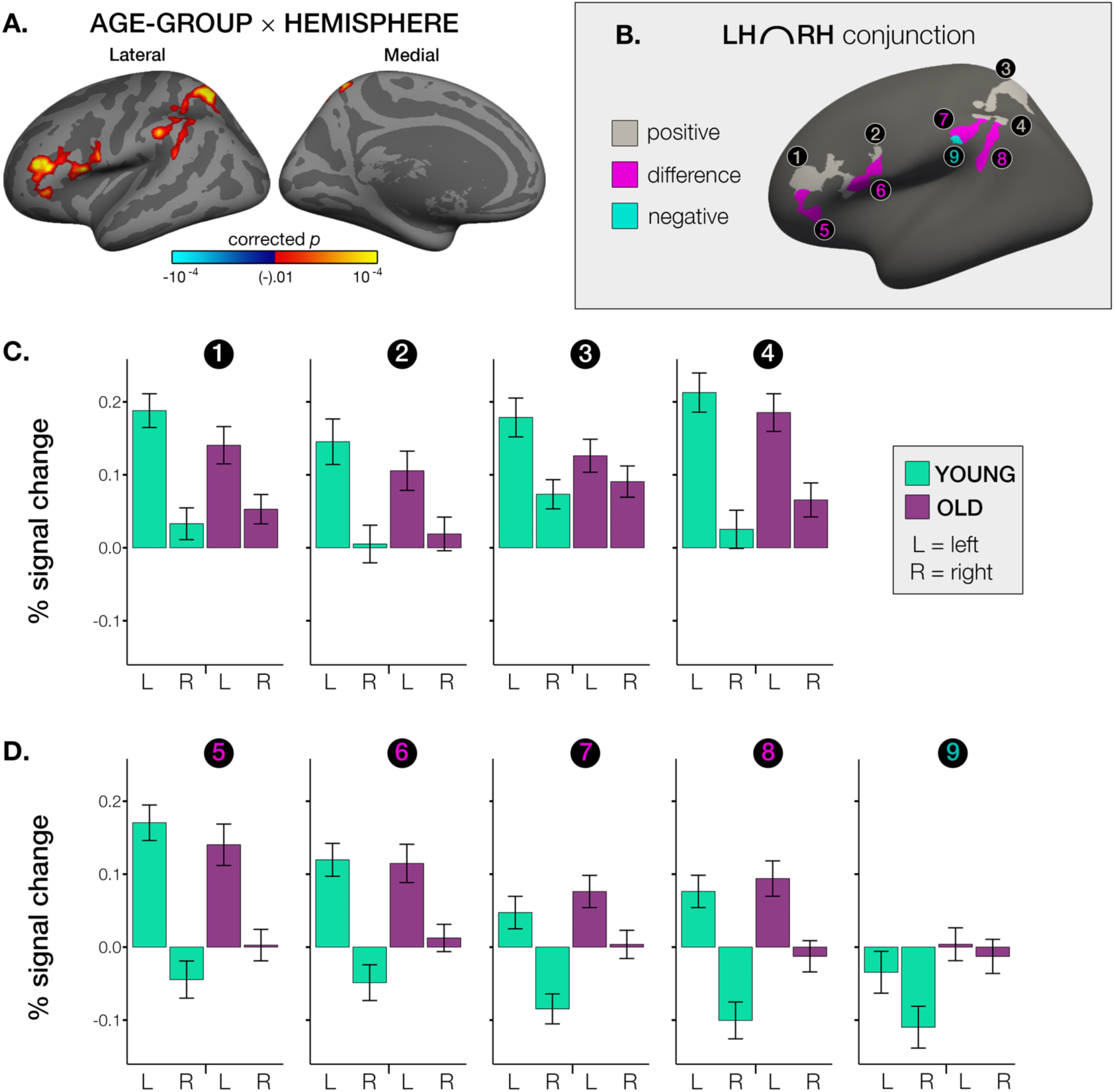
**A.** Cortical regions showing a significant Age-Group × Hemisphere interaction (clusters surviving FWE multiple comparison correction; vertex-wise *p* < 0.01; cluster-based *p* < 0.05). **B.** Resulting clusters were split according to average interhemispheric activation profile (Positive = positive effect in both hemispheres; Negative = negative effect in both hemispheres; Different = positive effect in one hemisphere and negative effect in the other) via interhemispheric conjunction analyses, resulting in 9 functionally distinct ROI’s (numbered 1-9 on the lateral surface). **C.** Activity plotted for each age-group and hemisphere in the 4 ROI’s showing positive effects in both hemispheres. **D.** Activity plotted for each age-group and hemisphere in the 4 ROI’s showing different effects in each hemisphere, and 1 ROI showing negative effects in both hemispheres. All group analyses were performed within the symmetrical space (the left symmetrical surface) which allowed direct homotopic comparison of activity between the hemispheres on a vertex-to-vertex basis. Error bars depict 95% confidence intervals. Note that plots are intended for evaluation of the vertex-wise interaction based on ROI-extracted data. Accordingly, effect sizes are likely inflated. LH = left hemisphere, RH = right hemisphere. See Tables 2, 3 and 4 for post-hoc descriptions of the effects.

In all 9 ROI’s, descriptive plots revealed that the interaction between Age-Group and Hemisphere pertained to markedly lower asymmetry in older adults relative to the young. We observed higher activation of the left –vs. right-hemisphere in all ROI’s (Fig. 4C,D). This difference between left and right activation magnitude was typically reflected by large effect sizes in both young and older adults, in all except one ROI (not in anterior supramarginal; see Table 2 for *left v. right* paired *t*-tests). Thus, functional asymmetry was nevertheless also present in older participants in these regions, but to a lesser extent.

**Table 2.** Asymmetry in ROI’s.

| Interhemispheric activation profile | ROI's | LEFT v RIGHT |  |  |  |
| --- | --- | --- | --- | --- | --- |
|  |  | YOUNG |  | OLD |  |
| | | LH-RH ($\bar{x}$ [SD]) | Effect (d') | LH-RH ($\bar{x}$ [SD]) | Effect (d') |
| Positive | rostral middle frontal | 0.16 (0.08)* | 1.8 | 0.09 (0.07)* | 1.3 |
|  | precentral | 0.14 (0.14)* | 1.0 | 0.09 (0.11)* | 0.8 |
|  | superior parietal | 0.11 (0.09)* | 1.2 | 0.04 (0.06)* | 0.6 |
|  | superior supramarginal | 0.19 (0.14)* | 1.4 | 0.12 (0.09)* | 1.3 |
| Different | pars triangularis | 0.21 (0.12)* | 1.7 | 0.14 (0.10)* | 1.4 |
|  | pars opercularis | 0.17 (0.10)* | 1.6 | 0.10 (0.09)* | 1.1 |
|  | supramarginal | 0.13 (0.11)* | 1.2 | 0.07 (0.08)* | 0.9 |
|  | supramarginal | 0.18 (0.13)* | 1.4 | 0.11 (0.11)* | 1.0 |
| Negative | anterior supramarginal | 0.08 (0.14)* | 0.5 | 0.02 (0.11) | 0.2 |

Post-hoc descriptions of asymmetry in regions of interest (ROI’s) identified via the whole-cortical Age-Group × Hemisphere interaction. Descriptives for the difference in hemispheric activation (left – right) and *significance results for *left v. right* paired *t*-tests (Bonferroni corrected at *p* < [0.05/18] 0.003) and corresponding effect sizes (Cohen’s d’) within young (columns 3:4) and older (columns 5:6) adults. ROI’s are ordered by interhemispheric activation profile (Positive = positive effect in both hemispheres; Negative = negative effect in both hemispheres; Different = positive effect in one hemisphere and negative effect in the other). Note that the outcome of this is a post-hoc description of the patterns underlying the vertex-wise interaction based on ROI-extracted data. Accordingly, effect sizes are likely inflated. LH = left hemisphere, RH = right hemisphere.

Of the four ROI’s that showed differing activation profiles depending upon hemisphere, plots indicated that the asymmetry pertained to positive memory effects in LH (in young and old) together with negative memory effects in the homotopic RH site in young participants, but not in older participants (Fig. 4D). Post-hoc one sample *t*-tests confirmed that within these regions, negative RH effects were significant in all cases in the young, and in all cases not significant in older participants (see Table 4). Further post-hoc *Old v. Young t-*tests across all 9 ROI’s revealed that older participants showed significantly greater RH effects only in regions exhibiting significant negative effects in the young (see Table 4; right columns). These age-related differences were typically reflected by medium and large effect sizes (though caution is warranted when interpreting effect size from ROI-extracted data; Kriegeskorte et al. 2010). Importantly then, we observed no significant age-related increase in contralateral RH activation in regions exhibiting positive effects in both hemispheres. Only one of these regions exhibited a trend towards higher recruitment in older adults (post-Bonferonni correction; superior supramarginal). Descriptive plots and post-hoc *Old v. Young t-*tests indicated that lower asymmetry in these regions was primarily driven by an apparent age-related decrease in LH memory effects (see Table 3; right columns). This was also substantiated by larger effect sizes for the age-group contrast in the LH (typically medium negative effect sizes) relative to the contralateral RH (typically low positive effect sizes). Together, these results indicate that all apparent age-related increases in contralateral RH recruitment pertained exclusively to regions that exhibited significant negative memory effects in the young, but showed no significant memory effects in older participants. Thus, lower asymmetry in older adults during retrieval-success was driven primarily either by less activity in the dominant left hemisphere, or a lack of negative memory effects in the contralateral right hemisphere.

**Table 3.** Left ROI activation and age-related differences.

| Interhemispheric activation profile | ROI's | Left |  |  |  | OLD v YOUNG |  |  |
| --- | --- | --- | --- | --- | --- | --- | --- | --- |
|  |  | YOUNG (T) | Sig | OLD (T) | Sig | Left (T [p]) | Sig | Effect (d') |
| Positive | rostral middle frontal | 15.9 | * | 10.8 | * | -2.7 (7.6e<sup>-3</sup>) | ° | -0.4 |
|  | precentral | 9.1 | * | 7.7 | * | -1.9 (6.5e<sup>-2</sup>) |  | -0.3 |
|  | superior parietal | 13.2 | * | 10.9 | * | -2.9 (4.3e<sup>-3</sup>) | * | -0.5 |
|  | superior supramarginal | 15.5 | * | 14.1 | * | -1.4 (0.2) |  | -0.2 |
| Different | pars triangularis | 13.7 | * | 9.7 | * | -1.6 (0.1) |  | -0.2 |
|  | pars opercularis | 10.4 | * | 8.6 | * | -0.3 (0.8) |  | -0.0 |
|  | supramarginal | 4.2 | * | 6.8 | * | 1.8 (7.5e<sup>-2</sup>) |  | 0.3 |
|  | supramarginal | 6.8 | * | 7.6 | * | 1.1 (0.3) |  | 0.2 |
| Negative | anterior supramarginal | -2.4 | ° | 0.3 |  | 2.0 (5.0e<sup>-2</sup>) | ° | 0.3 |

Post-hoc descriptions of left hemisphere effects underlying the whole-cortical Age-Group × Hemisphere interaction. Columns 3-6: Results from one sample *t*-tests in left hemisphere ROI’s for each age-group, ^O^pre- and *post-Bonferroni correction *p* < [0.05/18] 0.003. Column 7:9: Results from *Old v. Young* post-hoc *t*-tests of left hemisphere activity, ^O^pre- and *post-Bonferroni correction *p* < [0.05/9] 0.006 and corresponding effect sizes (Cohen’s d’). Note that in ROI’s exhibiting a positive interhemispheric activation profile, lower left hemisphere activity was most apparent in older adults. ROI’s are ordered by interhemispheric activation profile (Positive = positive effect in both hemispheres; Negative = negative effect in both hemispheres; Different = positive effect in one hemisphere and negative effect in the other). Note that the outcome of this is a post-hoc description of the patterns underlying the vertex-wise interaction based on ROI-extracted data. Accordingly, effect sizes are likely inflated. Sig = significance.

**Table 4.** Contralateral right ROI activation and age-related differences.

| Interhemispheric activation profile | ROI's | Right |  |  |  | OLD v YOUNG |  |  |
| --- | --- | --- | --- | --- | --- | --- | --- | --- |
| | | YOUNG (T) | Sig | OLD (T) | Sig | Right (T [p]) | Sig | Effect ($d'$) |
| Positive | rostral middle frontal | 3.0 | <sup>o</sup> | 5.1 | * | 1.3 (0.2) |  | 0.2 |
|  | precentral | 0.4 |  | 1.6 |  | 0.8 (0.4) |  | 0.1 |
|  | superior parietal | 7.2 | * | 8.3 | * | 1.2 (0.2) |  | 0.2 |
|  | superior supramarginal | 1.9 |  | 5.5 | * | 2.2 (2.7e<sup>-2</sup>) | <sup>o</sup> | 0.3 |
| Different | pars triangularis | -3.4 | * | 0.2 |  | 2.7 (7.1e<sup>-3</sup>) | <sup>o</sup> | 0.4 |
|  | pars opercularis | -3.9 | * | 1.3 |  | 3.8 (2.1e<sup>-4</sup>) | * | 0.6 |
|  | supramarginal | -8.1 | * | 0.4 |  | 6.1 (8.7e<sup>-9</sup>) | * | 0.9 |
|  | supramarginal | -7.8 | * | -1.2 |  | 5.1 (9.3e<sup>-7</sup>) | * | 0.8 |
| Negative | anterior supramarginal | -7.5 | * | -1.1 |  | 5.0 (1.2e<sup>-6</sup>) | * | 0.8 |

Post-hoc descriptions of contralateral right hemisphere effects underlying the whole-cortical Age-Group × Hemisphere interaction. Columns 3-6: Results from one sample *t*-tests in contralateral right hemisphere ROI’s for each age-group, ^O^pre- and *post-Bonferroni correction *p* < [0.05/18] 0.003. Columns 7:8: Results from *Old v. Young* post-hoc *t*-tests of activity in the contralateral right hemisphere, ^O^pre- and *post-Bonferroni correction *p* < [0.05/9] 0.006. Note that regions showing significantly greater contralateral activation in older adults were reliably associated with negative effects in young but not older adults. ROI’s are ordered by interhemispheric activation profile (Positive = positive effect in both hemispheres; Negative = negative effect in both hemispheres; Different = positive effect in one hemisphere and negative effect in the other). Note that the outcome of this is a post-hoc description of the patterns underlying the vertex-wise interaction based on ROI-extracted data. Accordingly, effect sizes are likely inflated. Sig = significance.

In agreement with ROI results, vertex-wise cortical analyses revealed that successful retrieval was associated with large-scale left-asymmetric activation across the cortex, irrespective of age (main effect of Hemisphere; Fig. 5, left column). Only one region in lateral occipital cortex showed rightward asymmetry irrespective of age-group. For further illustration of overall age-group similarity, we conducted a whole-cortical paired *t*-test in each age-group. This analysis revealed one cluster in transverse temporal cortex that exhibited rightward asymmetry in the young sample only. More generally, however, comparable left-dominant functional asymmetry was observed in both young and old age-groups—across the cortex (Fig. 5, middle and right panels, respectively). Further, a DSC of 0.87 substantiated the overlap in cortical asymmetry maps between age-groups.

**Figure 5.**
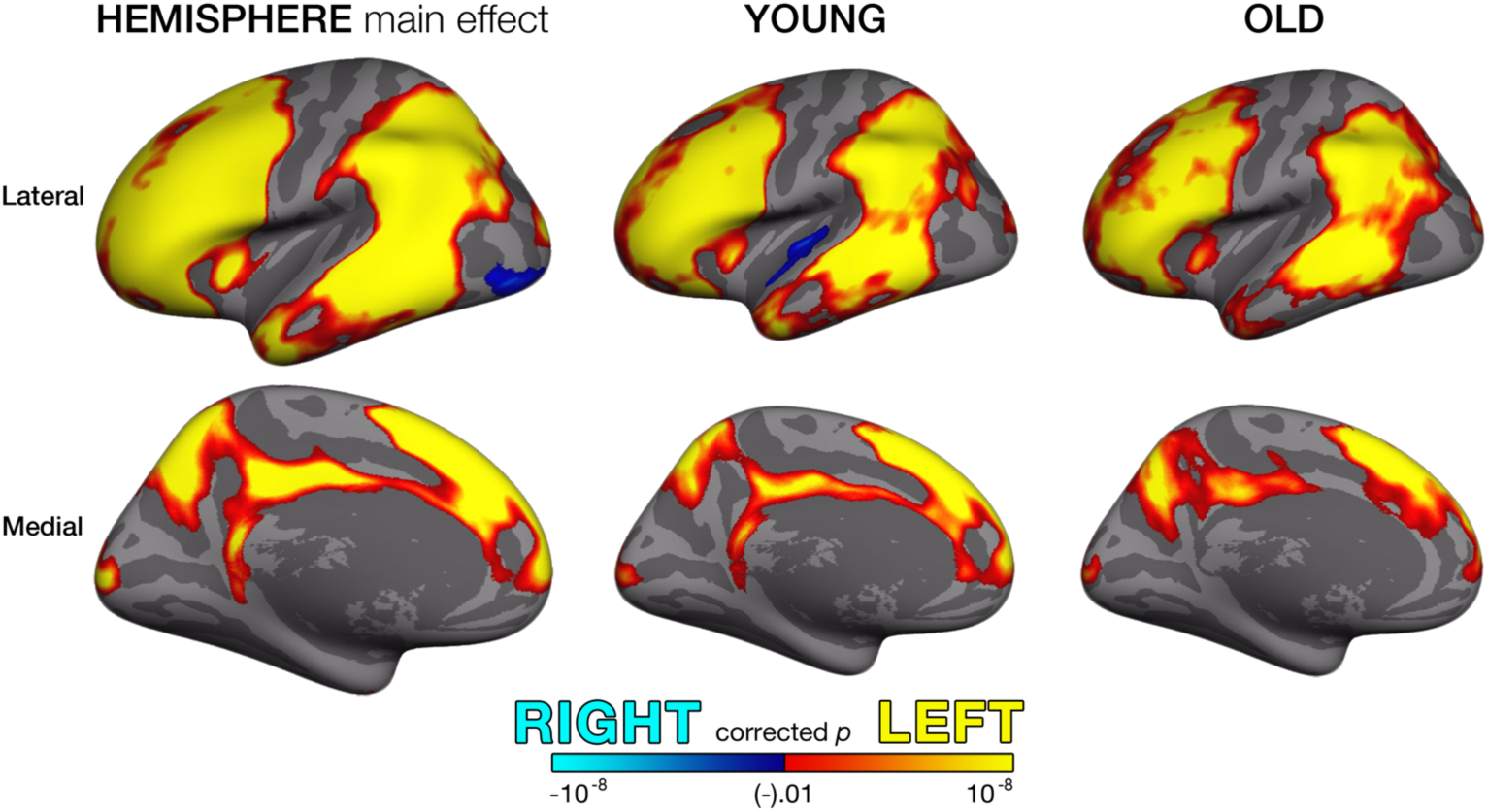
Cortical asymmetry maps. Warm colours represent regions where homotopic activity was significantly greater in the left hemisphere, whereas cold colours represent regions where homotopic activity was significantly greater in the right hemisphere. Results are shown for the main effect of Hemisphere (irrespective of age-group; left panel) and *left v. right* paired *t*-tests performed separately in young (middle panel) and older (right panel) age-groups. Note the extensive left-dominant functional asymmetry and high consistency between age-groups. All group analyses were performed within the symmetrical space (the left symmetrical surface) which allowed direct homotopic comparison of activity between the hemispheres on a vertex-to-vertex basis. All clusters survived FWE multiple comparison correction; vertex-wise *p* < 0.01; cluster-based *p* < 0.05.

Further vertex-wise analyses revealed a group difference in retrieval-success activity—irrespective of hemisphere—in a large cluster in supramarginal gyrus, extending along inferior postcentral and precentral gyrus into the anterior insula (main effect of Age-Group; Fig. 6). Although activity was significantly greater in these regions in older adults, descriptive plots confirmed that the difference pertained to a significant negative memory effect in younger participants not seen in older participants (see also Fig. 3). We reproduced all effects reported thus far after applying varying levels of surface-based smoothing. Results proved robust across smoothing levels (see Supplementary Fig. 2 for ANOVA-effects; see Supplementary Fig. 3 for conventional group-effects performed separately by hemisphere).

**Figure 6.**
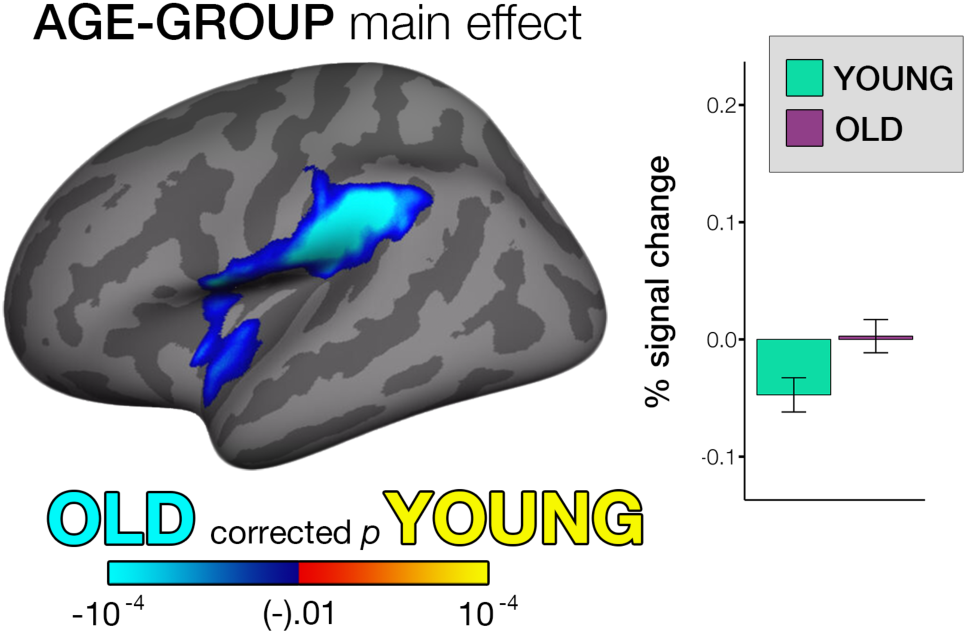
Main effect of Age-Group and plotted effects irrespective of hemisphere in young and older adults. Cold colours represent regions where activity was significantly greater in older adults, irrespective of hemisphere. All group analyses were performed within the symmetrical space (the left symmetrical surface) which allowed direct homotopic comparison of activity between the hemispheres on a vertex-to-vertex basis. All clusters survived FWE multiple comparison correction; vertex-wise *p* < 0.01; cluster-based *p* < 0.05. Error bars depict 95% confidence intervals.

Finally, we tested whether RH regions exhibiting retrieval-success effects nevertheless showed a left-dominant relative asymmetry. Here, conjunction analyses revealed that 59% of the vertices exhibiting significant activity in RH nevertheless exhibited significantly greater activity in the LH. Thus, left-dominant asymmetry was evident in a large proportion of the cortical network identified as active during retrieval-success in the present study— also where RH activation was significant (Fig. 7).

**Figure 7.**
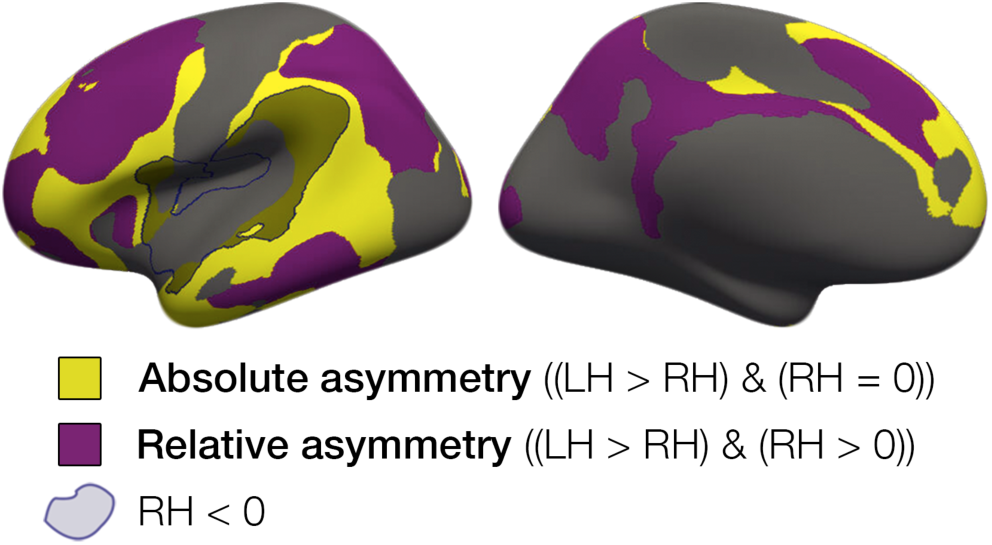
Cortical regions exhibiting absolute asymmetry (yellow; only significant in left) and relative asymmetry (purple; significant bilateral activation but significantly greater in left), and negative right effects (blue outline) on average across age-groups. All clusters survived FWE multiple comparison correction; vertex-wise *p* < 0.01; cluster-based *p* < 0.05. LH = left hemisphere, RH = right hemisphere.

### 3.4 Testing asymmetry relationships with memory and longitudinal memory change

We then tested the relationship between asymmetry and memory performance, and whether asymmetry-memory relationships differed between young and older adults. None of the tests (ANCOVA’s; age, sex controlled) in any of the 9 ROI’s revealed any significant main effect of Source memory performance upon asymmetry (all F[1,163] < 2.64, all *p* > .11 [uncorrected], all r^2^ < .01), nor any significant Age-group × Source memory interactions (all F[1,163] < 2.09, all *p* > .15 [uncorrected], all r^2^ < .01). Post-hoc analyses to determine whether asymmetry-memory relationships differed between old-low and old-high performance groups revealed no significant Group × Source memory interactions in any of the 9 ROI’s (all F[1,74] < 0.12, all *p* > .12; all r^2^ < .03). See Supplementary Tables 3 and 4 for ANCOVA analysis statistics.

Next, in a subset of our older participants (N=54) we used CVLT scores to estimate the extent of longitudinal memory change exhibited across the preceding ∼8 years (see Supplementary Fig. 1). CVLT delayed recall at Tp3 and memory performance in the fMRI task (conducted at Tp3) were highly correlated in the complete sample (r = .52, *p* = 10^-12^; see Supplementary Fig. 1A) and the older longitudinal subsample (r = .43, *p* = 10^-3^). We observed no significant relationships between asymmetry and longitudinal memory change in any of the 9 ROI’s found to exhibit age-related asymmetry differences (all *p* > .24 [uncorrected]). Further, no relationships between asymmetry and longitudinal memory change were found when using CVLT normative scores, i.e. T scores according to age and sex.

We additionally tested the relationship between memory performance and activation in the contralateral RH only. Again, we observed no significant relationships in any ROI for any of the above tests (all *p* > .18 [uncorrected]).

Thus, out of a total 90 tests probing brain-behaviour relationships (and group interactions) with current memory performance or longitudinal memory change, none revealed any significant relationships using a liberal statistical threshold even prior to multiple comparison correction (α = .05). We also tested whether recognition-reaction time (RT) on our fMRI task—as an indicator for processing efficiency—related to asymmetry, and found no significant asymmetry-RT relationships (see Supplementary Information). Finally, a posteriori power sensitivity analyses revealed that medium population effect sizes (13% explained variance) could be excluded given our sample size (power = .80; α = .05; two-tailed testing).

## 4. Discussion

We investigated whether functional brain asymmetry during successful memory retrieval is lower in older relative to younger adults, using methods to test asymmetry across the entire cortex. Our study yielded three main findings. First, the analyses revealed lower asymmetry in older adults in widespread prefrontal and parietal regions. However, we observed no evidence that lower asymmetry corresponded to an apparent age-related increase in recruitment of the contralateral right hemisphere in older adults. Rather, contralateral activations that were significantly greater in older adults pertained exclusively to regions that reliably signalled retrieval failure in the young (i.e. exhibited negative memory success effects), but that showed no significant effects in older adults. Results thus indicated that the pattern of lower asymmetry in older adults was primarily driven by either reduced deactivation in right contralateral cortex, or by less retrieval activity in the dominant left hemisphere. Second, we found direct evidence that activation of the cerebral cortex during successful retrieval may be characterised by an extensive left-dominant functional asymmetry, encompassing a large proportion of the cortical retrieval network identified as active here. This result proved to be highly reproducible in both age-groups. Hence, despite the apparent age-related reduction in asymmetry, our results indicate that the older brain relies on processing strategies that are largely asymmetric during retrieval—similar to younger adults—and that asymmetry facilitated successful retrieval in both age-groups. Third, we found no evidence that individual differences in the magnitude of asymmetry were related to current memory performance, nor the extent of longitudinal memory change exhibited across an ∼8 year period prior to scanning in older adults.

### 4.1 Age-related differences in asymmetry

In line with previous research, our results indicate that lower prefrontal asymmetry may be a feature of the older brain during memory retrieval (Bäckman et al. 1997; Cabeza et al. 1997, 2002, 2004; Madden et al. 1999; Grady et al. 2002; Rossi et al. 2004; Berlingeri et al. 2013; Morcom and Johnson 2015). Yet our findings also extend the previous literature. Firstly, we observed that lower asymmetry in older age may be a more widespread phenomenon across the cortex when recalling a memory, occurring across large swathes of parietal cortex where retrieval-induced activation is often evident (Wagner et al. 2005; Berlingeri et al. 2013; Frithsen and Miller 2014). Of importance, however, we observed that lower asymmetry was manifest in a different pattern than what is typically reported (Cabeza 2002; Cabeza and Dennis 2012). Namely, we found no evidence to indicate that older adults exhibit significantly greater activation of the contralateral hemisphere than younger adults—or show a shift towards bilaterality—during retrieval. Rather, in frontal and parietal regions exhibiting positive memory effects in both hemispheres (hereafter termed contralateral co-activation), the data indicated a reduction in responsivity of the dominant left hemisphere in older adults, combined with no significant age-related change in right homotopic cortex. Thus, our study yielded no convincing evidence to indicate that older adults recruit additional resources from the contralateral hemisphere; the pattern arguably emphasised within a compensation framework of HAROLD, given consensus that the term typically describes a situation in which brain activity is greater in older adults (Cabeza et al. 2018). Previously, additional contralateral recruitment in older adults has been proposed to account for insufficient engagement of the hemisphere more specialised to perform the task (Cabeza et al. 1997, 2004; Cabeza 2002; Persson and Nyberg 2006; Cabeza and Dennis 2012). Since we observed primarily lower activation of the dominant left hemisphere in these regions, our results are inconsistent with the compensation view of HAROLD during retrieval.

Secondly, we found that all regions that did show significantly greater contralateral activity in older adults were reliably associated with negative memory effects in young participants (i.e. greater activation during retrieval failure). In these, homotopic regions exhibited a functional dissociation between hemispheres, such that positive left activation was associated with contralateral right deactivation in the young, but neither activation nor deactivation of contralateral cortex in the old. These included two clusters located in inferior frontal cortex and two in supramarginal gyrus, and appeared to broadly correspond to regions typically encompassed within either the so-called default network (Buckner et al. 2008; Yeo et al. 2011) or the ventral network associated with stimulus-driven attentional capture (Corbetta et al. 2008; Yeo et al. 2011). The lack of negative memory effects in older adults suggests that a similar level of regional neural activity was present in trials recalled successfully and trials associated with retrieval failure. Thus, the present results may complement previous reports claiming that aging may be associated with a reduction in the ability to suspend neural processing not conducive to current cognitive goals (Lustig et al. 2003; Grady et al. 2006; Maillet and Rajah 2014). For example, one can speculate that given its established role in attention reorienting (Corbetta and Shulman 2002; Corbetta et al. 2008), the suppression of activity in right supramarginal cortex may be facilitative to memory goals, yet older adults showed an apparent deficit in this ability. Relatedly, cortical regions we identified showing no contralateral deactivation in older adults could reflect reduced functional inhibition of homotopic cortex, potentially as a consequence of age-related degeneration of callosal fibers (Buckner and Louis 2004; Head et al. 2004; Persson et al. 2006; Sullivan and Pfefferbaum 2006; Hou and Pakkenberg 2012; Yeatman et al. 2014) that may affect the balance in the distribution of hemispheric processing (Szczepanski and Kastner 2013). Nevertheless, the pattern we observed in cortical regions exhibiting contralateral co-activation does not support a disinhibition account, because we observed no significant age-related increase in contralateral activation. Since there is evidence the corpus callosum mediates both contralateral excitation and inhibition in a manner that may be regionally dependent (van der Knaap and van der Ham 2011; Roland et al. 2017), future research should assess the longitudinal link between regional callosal atrophy and regional changes in asymmetry. This may shed light on mechanisms mediating hemispheric distribution of neural resources, and potential alterations in this ability during aging.

The pattern of lower asymmetry in older adults observed here primarily reflected either a lack of negative memory effects in right contralateral cortex, or lower activation of the dominant left hemisphere. Overall this suggests that age-related deficits in the brain activity associated with memory retrieval can drive lower asymmetry in older brains. This is in line with the well-established view that aging is associated with the de-differentiation of previously specialised neural systems (Park et al. 2004, 2012; Park and McDonough 2013; Koen et al. 2019), and may also reconcile well with studies demonstrating that the regionally-dependent dynamic range of activity available to older adults may be lower across varying task conditions (Garrett et al. 2011, 2013; Kennedy et al. 2017; Amlien et al. 2018), a potential alternative account that may explain findings of lower-to-absent negative memory effects in older adults (Mattson et al. 2014; de Chastelaine et al. 2015). In particular, our finding that absent negative effects explained all instances of apparent over-activation in older adults agrees well with a similar report during retrieval (Morcom et al. 2007). Conversely, our results are difficult to reconcile with previous cross-sectional findings indicating higher contralateral activation in older adults during retrieval (Bäckman et al. 1997; Cabeza et al. 1997, 2004; Madden et al. 1999; Cabeza 2002; Rossi et al. 2004; Bishop et al. 2010; Cabeza and Dennis 2012; Berlingeri et al. 2013; Morcom and Johnson 2015)—most frequently interpreted as evidence for neural compensation in aging. Although the reasons for the inconsistent results between these and the present study are almost certainly multi-faceted—potentially spanning differences between imaging modalities (PET/fMRI), task type and the type of memory being probed among others— differences in experimental design may constitute a particularly important factor. For example, whether memory activity is contrasted against a task-related (Cabeza et al. 2002) or an implicit baseline condition (Bäckman et al. 1997; Cabeza et al. 1997; Madden et al. 1999) may have marked implications for the interpretation of results, as only the former allows one to isolate brain regions specific to retrieval-success from regions recruited during both successful and unsuccessful retrieval attempts. Differences between task-related contrasts and the cognitive processes they isolate could also account for some inconsistent results. Our results also only partially agree with those of Berlingeri et al. (2013); during simple old/new recognition, their results indicate that some frontal and parietal patterns of lower asymmetry may have been driven by lower dominant-hemisphere activation (the right in their study), whereas contralateral over-activation appeared most apparent in driving the patterns in others. However, in our large cross-sectional sample, we were unable to detect convincing evidence for the presence of apparent age-related contralateral over-activation during retrieval-success, except in the context of absent negative memory effects in older adults. Extant evidence suggests that lower asymmetry may reflect a general phenomenon in aging across tasks—at least in prefrontal cortex (Bäckman et al. 1997; Cabeza et al. 1997, 2004; Madden et al. 1999; Rosen et al. 2002; Cabeza 2002; Logan et al. 2002; Morcom et al. 2003, 2007; Bergerbest et al. 2009; Duverne et al. 2009; Reuter-Lorenz and Park 2010a; Morcom and Friston 2012; Davis et al. 2012; Berlingeri et al. 2013). Hence, we emphasise that our results only apply to the domain of memory retrieval, also because age-related asymmetry changes are likely to be task-specific (Berlingeri et al. 2013). Indeed, tasks employing parametric load manipulations provide evidence that increased bilateral recruitment may parallel increases in task-demand in both younger and older adults, but that older adults recruit contralateral resources at comparatively lower loads (Reuter-Lorenz and Cappell 2008; Reuter-lorenz and Park 2010; Schneider-Garces et al. 2010), possibly explaining age-related asymmetry differences across varying tasks (Berlingeri et al. 2013). However, that older adults in the present study were significantly impaired in source memory performance also suggests they were more cognitively taxed, possibly suggesting our results during memory retrieval may also not align well with alternative theoretical accounts (Compensation-related Utilization of Neural Circuits Hypothesis; Reuter-lorenz and Stanczak 2000). Studies employing retrieval-related load manipulations are needed to elucidate this. Nevertheless, the present results do not exclude the presence of compensatory activity in the ipsilateral hemisphere, nor in contralateral regions that are non-homotopic. Still, our results may call into question the focus on higher contralateral brain responses in older adults during memory retrieval (Cabeza et al. 2002, 2018; Grady 2012), highlighting instead that lower asymmetry does not imply a higher contralateral brain response, and that patterns driving lower asymmetry in older brains may reflect age-related neural deficits.

### 4.2 Asymmetry: memory performance and memory change

The degree of asymmetry was not related to current memory performance. Thus, none of the accepted premises for compensation were found for the apparent asymmetry reduction observed here (Cabeza et al. 2018): we observed no age-related contralateral over-activation, and lower asymmetry conferred no beneficial effect on memory performance. It has recently been proposed that lower asymmetry may rather correlate with performance on a trial-by-trial basis within individuals, as opposed to memory performance compared across individuals (Wang and Cabeza 2017). Although the present study was not designed to assess intra-individual compensation, this would arguably still be dependent on evidence that older adults benefit more from bilateral activation than the young (across trials) if contralateral compensation is an important feature of the aging brain during retrieval. However, our results indicated that asymmetry was associated with retrieval-success in both age-groups. Thus, at the group level, both young and older adults benefited from the asymmetric recruitment of cortical circuits, indicating that both young and older brains relied on similar neural processes during retrieval.

The magnitude of asymmetry was also unrelated to the extent of longitudinal memory decline exhibited across the preceding ∼8 years in a subset of 54 older participants. Thus, we found no evidence that retrieval-induced asymmetry is a marker for age-related cognitive preservation or decline. Given that research is conflicting regarding the cognitive outcomes of lower frontal asymmetry (Eyler et al. 2011), and that an age-related difference in activity is not synonymous with a change in activity occurring with age (Rugg 2017), we note that future large-scale longitudinal studies are needed to test the relationship between changes in asymmetry over time, and memory maintenance or decline in aging individuals.

### 4.3 Asymmetry common to young and old

We observed direct evidence that the human cerebral cortex may rely disproportionately on memory signals localised in the left hemisphere during memory retrieval, and that this asymmetry potentially occurs on an unprecedented scale across the cortex. Moreover, this left-dominant asymmetry showed high consistency between young and old age-groups. This is in line with many newer reports indicating an age-invariant left-dominant asymmetry during retrieval (Morcom et al. 2007; Duverne et al. 2008; Angel et al. 2013; de Chastelaine et al. 2016; Wang et al. 2016). Hence, we emphasise that we observed extensive asymmetry also in older brains. Indeed, older adults exhibited strong asymmetry even in regions characterised by an apparent age-related asymmetry reduction. Thus, we found only relative differences in hemispheric asymmetry between age-groups. Furthermore, lower asymmetry was evident also in regions exhibiting significant bilateral activation common to both age-groups, and we observed not only few qualitative differences in the activation maps of young and old adults to indicate asymmetry differences (see Fig. 3), but also a high degree of activation overlap between age-groups in each hemisphere. Taken together, we argue that these observations have implications for future research concerned with identifying asymmetry changes in aging. Adopting similar methodologies that do not restrict the search for asymmetry to specific brain regions may promote a deeper understanding of processing strategies in the cerebral hemispheres and their relation to aging.

### 4.4 Overarching considerations

Generally, asymmetries have been reported in frontal and parietal regions during memory retrieval (Kalpouzos and Nyberg 2010), and thus our results contribute direct evidence that retrieval-induced asymmetry may occur on a more global scale than previously identified. Strengthening the potential generalizability of these findings, the activation patterns we observed delineating retrieval-success separately by hemisphere were largely in agreement with recent research (McDermott et al. 2009; Spaniol et al. 2009; Huijbers et al. 2013; Kim 2016). However, the extensive left-dominant asymmetry observed here also stands in stark contrast to earlier research indicating that retrieval is more dependent on right prefrontal cortex (Tulving et al. 1994; Cabeza et al. 1997; Cabeza and Nyberg 2000), an observation that formed part of the hemispheric encoding/retrieval asymmetry model (Tulving et al. 1994; Nyberg et al. 1996, 1998). While the reasons for this discrepancy remain elusive, evidence from a multi-study PET investigation suggests that processes that support retrieval only indirectly may account for some previous reports of right-prefrontal asymmetry during retrieval (Lepage et al. 2000). Nevertheless, this does not reconcile well with behavioural investigations of retrieval-asymmetry (Rossi et al. 2001, 2004), nor with whole-brain assessments that intriguingly reveal right-dominant asymmetry during picture recognition (Berlingeri et al. 2013). Alternatively, fMRI (Rugg et al. 1999; Slotnick et al. 2003; Dobbins and Wagner 2005) and patient data (Duarte et al. 2005) suggests that left frontal cortex is critical for the retrieval of contextual information, whereas right frontal cortex may support recognition/familiarity-based retrieval in the absence of recollection (Nolde et al. 1998; Dobbins et al. 2004), which may go some way to resolving inconsistencies in reported retrieval-asymmetries. Indeed, although our results argue against right-prefrontal dominance (Cabeza et al. 1997; Habib et al. 2003), we also found that large portions of frontal cortex were characterised by only relative asymmetry (significant bilateral effects but significantly greater in the left), thus also evidencing extensive right frontal recruitment during retrieval-success. Regardless, our results seem consistent with a role for left prefrontal dominance during the retrieval of memories with concurrent contextual information (Duarte et al. 2005).

We note that because our paradigm probed retrieval using visual and auditory probes, we chose to always contrast within-subject activity elicited during successful versus unsuccessful retrieval, to control for asymmetries potentially arising from the sensory conditions during testing. This task-related baseline also allowed us to test asymmetry-differences in activity specific to successful retrieval, and avoid confounds associated with age-related differences in raw BOLD activity (Lustig et al. 2003; Andrews-Hanna et al. 2007; Fjell et al. 2014). Although a similar rationale led to our use of source memory versus miss as the fMRI contrast, a notable limitation arises from this decision: if some trials classified as source memories (widely acknowledged to compare with recollection; Yonelinas 2001; Davachi et al. 2003; Diana et al. 2007; Wixted and Squire 2011) also included the neural correlates associated with more familiarity-based or less-confident memory judgements, our chosen contrast would not be able to separate these forms of memory processing. Hence, the activity captured here, and associated age-related differences identified therein, is not specific to source memory success. This is true even though our paradigm also enabled categorising item-only memories in order to regress their influence out of the fMRI timeseries. Nevertheless, because the present study aimed to identify age-effects in retrieval-asymmetry, our fMRI contrast—which undoubtedly captures retrieval-success activity (Morcom et al. 2007; Duverne et al. 2008; Wang and Cabeza 2017)—is arguably ideally suited to this goal. That is, to contrast retrieval activity between groups that differ in memory ability, we chose to restrict analyses to trials for which an objective threshold-level of memory had been attained (i.e. source memory). Of principal importance, we also aimed to avoid a scenario in which age-related differences in activity may be more ascribable to differences in a baseline condition that may in and of itself exhibit age-related differences (McDonough et al. 2013; Mitchell et al. 2013; de Chastelaine et al. 2016) but may only be indirectly related to retrieval-success (Henson et al. 2000; Kim 2010). With this in mind, the authors opted not to use a source versus item-memory contrast, namely because its inverse (i.e. item > source or similar) is often adopted to identify neural correlates for post-retrieval monitoring processes thought to be more engaged during the retrieval of less-strong memory representations (Henson et al. 2000; Dobbins et al. 2004; Cabeza et al. 2008; Donaldson et al. 2010; Kim 2010; de Chastelaine et al. 2016), and age-related effects in post-retrieval monitoring have been found at both the behavioural (McDonough and Gallo 2013) and neural level (McDonough et al. 2013; Mitchell et al. 2013; de Chastelaine et al. 2016). Thus, our chosen contrast intended to avoid later interpretive issues potentially arising from introducing age-related confounds in the task-related baseline, although at the expense of perfectly isolating source memory-specific activity—for which a source versus item-memory contrast is doubtlessly advantageous (Davachi et al. 2003; Woodruff et al. 2005; Rugg and Vilberg 2013; King et al. 2015; de Chastelaine et al. 2016; Wang and Cabeza 2017).

Finally, because homotopic regions are not thought to be confounded by differences in neurovascular coupling (Miezin et al. 2000; Morcom and Johnson 2015), our whole-cortical asymmetry approach may help circumvent this common confound when comparing regional activity between age-groups (D’Esposito et al. 1999; West et al. 2019). Indeed, our use of a symmetrical surface template to perform hemispheric comparisons was informed by recent reports demonstrating both the utility of such an approach, and that such methods are gaining traction (Greve et al. 2013; Maingault et al. 2015; Takaya et al. 2015; Tobyne et al. 2016; Greve and Fischl 2017). However, it might be argued that this method may result in a degree of loss in anatomical specificity, potentially above what is apparent in standard templates modelled on hemisphere-specific landmarks. Nevertheless, we believe that the extent of the effects documented here and the high level of reproducibility in our asymmetry results between richly populated age-samples is encouraging, and together suggest that any anatomical specificity lost during the reregistration process likely had minimal impact on the overall results observed.

### 4.5 Conclusion

In conclusion, the present results confirm that lower functional asymmetry may be an important feature of the older brain during memory retrieval. Yet our study challenges common assumptions of a neural compensation account to HAROLD insofar as the model applies to memory retrieval, suggesting instead that apparent age-related asymmetry reduction may arise from less activation or less deactivation in older adults—a pattern more suggestive of age-related functional deficits—and that retrieval-induced asymmetry may be unrelated to either current memory performance, memory preservation or decline in neurocognitive aging. Future research should assess longitudinal alterations in brain asymmetry using unbiased methods to delineate asymmetry, and assess the relevance of this for understanding episodic memory change in the aging brain.

## Acknowledgements

We thank Roberto Cabeza for insightful discussions during the preparation of this manuscript. We are also indebted to Lars Nyberg for providing many extensive and invaluable comments on earlier drafts of this paper.

