## Supplementary Information for "Age-related differences in functional asymmetry during memory retrieval revisited: no evidence for contralateral over-activation or compensation"

**Supplementary Table 1**

Main demographic and neuropsychological variables

| Age-group | Sample descriptives | All | Old v Young (T [p]) | Effect (d') |
|---|---|---|---|---|
| Young | Sex (female:male) | 56 : 33 | - | |
| | Age | 26.8 (5.0) | - | |
| | Age Range | 18.5 : 38.9 | - | |
| | MMSE | 29.1 (1.0) | - | |
| | Vocabulary | 63.2 (7.0) | - | |
| | CVLT Learning | 58.6 (14.1) | - | |
| | CVLT 30' | 13.8 (2.6) | - | |
| Old | Sex (female:male) | 40 : 36 | - | |
| | Age | 67.9 (5.2) | - | |
| | Age Range | 60.2 : 80.8 | - | |
| | MMSE | 28.6 (1.3) | -2.6 ($9.6e^{-3}$)* | -0.4 |
| | Vocabulary | 65.6 (7.6) | 1.9 ($5.4e^{-2}$) | 0.3 |
| | CVLT Learning | 50 (10.7) | -4.3 ($2.8e^{-5}$)* | -0.7 |
| | CVLT 30' | 11.1 (2.7) | -6.2 ($4.1e^{-9}$)* | -1.0 |

Main sample demographics and raw neuropsychological scores (8 young participants lacked neuropsychological data). Where appropriate, results from *Old v. Young t*-tests are reported (*Bonferroni corrected at $p < [0.05/4]\ 0.01$) with corresponding effect sizes (Cohen's d'). Descriptive statistics represent $\bar{x}$ (SD), range or frequencies.

**Supplementary Table 2**

Cluster statistics for all cortical analyses

| Analysis | Contrast | Age-group | Hemisphere (**) | Max sig. | CWP | Size($mm^2$) | MNI(x,y,z) | Area |
|---|---|---|---|---|---|---|---|---|
| Conventional analysis (performed separately by hemisphere) | Source Memory v Miss | YOUNG | LH | 31.6 | 4 | 53990.7 | (-37, -61.7, 42.7) | lh inferiorparietal |
| | | | | -13.8 | 2.4 | 4830.9 | (-51.9, -28.7, 21.1) | lh supramarginal |
| | | | RH | 24.6 | 4 | 21409.5 | (-3.8, -67.8, 30.6) | rh precuneus |
| | | | | 17.2 | 2.8 | 4862.1 | (-33.5, 7.6, 55.1) | rh caudalmiddlefrontal |
| | | | | -14.2 | 3.2 | 8147.7 | (-57.2, -40.4, 33.2) | rh supramarginal |
| | | | | 11.2 | 1.6 | 965.6 | (-9.5, 14.5, 48.4) | rh superiorfrontal |
| | | | | 9.4 | 2.7 | 4148.1 | (-21.3, 60, 1.8) | rh rostralmiddlefrontal |
| | | | | -6.7 | 1.4 | 1217.6 | (-23.3, 47.6, 26) | rh rostralmiddlefrontal |
| | | OLD | LH | 27.4 | 4 | 62767.6 | (-5.5, -70, 44.6) | lh precuneus |
| | | | RH | 21.6 | 4 | 29279.5 | (-5.7, -68.9, 45.6) | rh precuneus |
| | | | | 17.0 | 2.9 | 12037.3 | (-31, 1.6, 54.2) | rh caudalmiddlefrontal |
| Asymmetry analysis: 2 (Age-Group) x 2 (Hemisphere) ANOVA | Age-Group × Hemisphere | YOUNG v OLD | LH v RH | 5.0 | 2.1 | 1218.6 | (-24.6, -60.6, 54.3) | superiorparietal |
| | | | | 4.6 | 2.4 | 1705.7 | (-38.4, 40.3, 13.6) | rostralmiddlefrontal |
| | | | | 3.9 | 1.8 | 853.1 | (-60.1, -30.8, 33.1) | supramarginal |
| | M.E. Hemisphere | YOUNG v OLD | LH v RH | 41.5 | 4 | 42774.7 | (-41.8, 38.7, 2.1) | rostralmiddlefrontal |
| | | | | 11.4 | 1.8 | 1419.7 | (-21, -96.8, 9.1) | lateraloccipital |
| | | | | -5.4 | 1.6 | 920.7 | (-42.9, -81.5, -10.9) | lateraloccipital |
| | M.E. Age-group | YOUNG v OLD | LH v RH | -5.7 | 2 | 3840.7 | (-51, -27.4, 19.5) | supramarginal |

| | | | | | | | |
|---|---|---|---|---|---|---|---|
| Left v Right | YOUNG | LH v RH | 24.6 | 4 | 38904.7 | (-39.4, 40.3, 3.8) | rostralmiddlefrontal |
| | | | 6.4 | 1.7 | 1149.2 | (-11.9, -102.2, 5.9) | lateraloccipital |
| | | | -5.6 | 1.5 | 589.9 | (-44, -23.1, -1.3) | transversetemporal |
| | OLD | LH v RH | 21.2 | 4 | 18031.6 | (-43, 38.2, -0.5) | parstriangularis |
| | | | 19.8 | 4 | 18011.3 | (-49, -40.7, -4.1) | bankssts |
| | | | 6.5 | 1.7 | 963.7 | (-22, -96.1, 9.6) | lateraloccipital |

Cluster statistics for all cortical analyses. All analyses pertained to the *source memory v miss* contrast (source memory success). First, fMRI analyses were performed separately by hemisphere in a conventional manner (top section). Then, asymmetry analyses were performed whereby whole-cortical activity was contrasted between homotopic hemispheres (bottom sections). (**) Note that cluster results from asymmetry analyses cannot be delineated by hemisphere (LH v RH). The reported clusters survived $p < 0.05$ FWE correction with a cluster defining threshold of $p < 0.01$. Cluster significance was tested through permutation testing. Max sig. = maximum voxel significance; CWP = cluster-wise probability (log transformed). See **Figure 3**, **Figure 4A**, **Figure 5** and **Figure 6** for corresponding visual illustrations.

**Supplementary Table 3**

ROI analysis (ANCOVA's)

| Interhemispheric activation profile | ROI's | Source memory | | Age-group × Source Memory | |
|---|---|---|---|---|---|
| | | F (p) | $r^2$ | F (p) | $r^2$ |
| Positive | rostral middle frontal | 2.64(0.11) | 0.01 | 0.21(0.64) | < 0.01 |
| | precentral | 0.38(0.54) | < 0.01 | 0(0.96) | < 0.01 |
| | superior parietal | 0.72(0.4) | < 0.01 | 0.42(0.52) | < 0.01 |
| | superior supramarginal | 1.43(0.23) | 0.01 | 0.4(0.53) | < 0.01 |
| Different | pars triangularis | 0.21(0.65) | < 0.01 | 0.04(0.83) | < 0.01 |
| | pars opercularis | 0.43(0.51) | < 0.01 | 0.25(0.62) | < 0.01 |
| | supramarginal | 0.52(0.47) | < 0.01 | 0.14(0.71) | < 0.01 |
| | supramarginal | 1.48(0.23) | 0.01 | 2.09(0.15) | 0.01 |
| Negative | anterior supramarginal | 0.03(0.86) | < 0.01 | 0.9(0.35) | 0.01 |

Results from ANCOVA’s (Age-group, Source memory performance, Age-group × Source memory as factors; age, sex controlled) testing asymmetry-memory relationships and whether asymmetry-memory relationships differed between young and older adults. No significant main effect of Source memory performance, nor Age-group × Source memory interaction was found upon asymmetry for any of the 9 ROI’s exhibiting age-related asymmetry differences.

**Supplementary Table 4**

ROI analysis (ANCOVA's)

| Interhemispheric activation profile | ROI's | performance-Group × Source Memory F (p) | $r^2$ |
|---|---|---|---|
| Positive | rostral middle frontal | 0.33(0.57) | < 0.01 |
| | precentral | 0.15(0.7) | < 0.01 |
| | superior parietal | 0.35(0.56) | < 0.01 |
| | superior supramarginal | 2.41(0.12) | 0.03 |
| Different | pars triangularis | 0.26(0.61) | < 0.01 |
| | pars opercularis | 0.13(0.72) | < 0.01 |
| | supramarginal | 0.76(0.39) | 0.01 |
| | supramarginal | 0.48(0.49) | 0.01 |
| Negative | anterior supramarginal | 0.14(0.71) | < 0.01 |

Results from ANCOVA's testing whether asymmetry-memory relationships differed between low performing and high performing older adults (performance-Group, Source memory performance, performance-Group × Source memory as factors; age, sex controlled). No significant performance-Group × Source memory interaction was found upon asymmetry for any of the 9 ROI's exhibiting age-related asymmetry differences.

***Supplementary analysis: testing asymmetry relationships with reaction time***

In line with a reviewer suggestion, we also tested the relationship between asymmetry and recognition reaction time (RT) on our fMRI task (correct response to Q1; see Figure 2), and whether asymmetry-RT relationships differed between young and older adults, or between old-low and old-high memory performance groups (similar to the main analyses). None of the tests (ANCOVA's; age, sex controlled) in any of the 9 ROI's revealed any significant main effect of RT upon asymmetry (all $F[1,163] < 2.68$, all $p > .10$ [uncorrected], all $r^2 < .01$). One of the 9 ROI's (superior supramarginal; cluster #4) was found to exhibit a significant Age-group × RT interaction ($F[1,163] < 4.49$, $p = .04$ [uncorrected], $r^2 = .03$), and the same ROI was found to exhibit a significant performance-Group (old-low/old-high) × RT interaction in older adults ($F[1,74] < 4.06$, $p = .05$ [uncorrected], $r^2 = .05$). Post-hoc testing of the underlying effects via linear regressions (age, sex corrected) between asymmetry and RT separately within each Age-group and older performance-Group revealed no significant asymmetry-RT relationships in neither young adults ($p = .07$ [uncorrected]), older adults ($p = .17$

[uncorrected]), nor old low- ($p = .14$ [uncorrected]) or high-performers ($p = .08$ [uncorrected]). Thus, we also observed no significant asymmetry-RT relationships in any the 9 ROI's exhibiting age-related asymmetry differences.

***Figure captions***

***Supplementary Figure 1.*** **A.** Scatterplot of the relationship between memory performance on the CVLT delayed recall task and source memory performance on our fMRI task in young and older adults. Memory performance on each was measured at timepoint 3 (Tp3). **B.** Longitudinal CVLT data spanning back ~8 years was used to estimate memory change over time. Data-points represent the per-participant slope (words per year across ~8 years) of the linear model of CVLT delayed recall against time, and are shown plotted against age at the final timepoint (Tp3). Shaded area around the line indicates SEM.

***Supplementary Figure 2.*** Main ANOVA results (uncorrected significance) reproduced applying varying levels of 2D surface-based smoothing (FWHM = 0mm, 2mm, 4mm, 6mm; row-wise denotations) to the functional data on the native cortical geometry of the individual prior to first-level statistical analyses. Compare with Figures 4, 5 and 6. Note that a slightly different upper visualization threshold is applied to the Hemisphere main effect here (compared to Fig. 5) to keep the visualization of smoothing-levels consistent across effects. All group analyses were performed within the symmetrical space (the left symmetrical surface) which allowed direct homotopic comparison of activity between the hemispheres on a vertex-to-vertex basis. Hemisphere = main effect of Hemisphere, Age-group = main effect of Age-group.

***Supplementary Figure 3.*** Results from conventional fMRI analyses (uncorrected significance) performed separately by hemisphere for young (panels 1 and 3) and older adults (panels 2 and 4). Note that the data from both left and right hemispheres is registered to the left symmetrical surface (left-most panels and right-most panels, respectively). Results were reproduced applying varying levels of 2D surface-based smoothing (FWHM = 0mm, 2mm, 4mm, 6mm; row-wise denotation) to the functional data on the native cortical geometry of the individual prior to first-level statistical analyses. For visualization, only lateral views are shown. Compare with Figure 3.
